# Dysfunction of grey matter NG2 glial cells affects neuronal plasticity and behavior

**DOI:** 10.1101/2021.08.20.457086

**Authors:** Aline Timmermann, Ronald Jabs, Anne Boehlen, Catia Domingos, Magdalena Skubal, Wenhui Huang, Frank Kirchhoff, Christian Henneberger, Andras Bilkei-Gorzo, Gerald Seifert, Christian Steinhäuser

**Affiliations:** Institute of Cellular Neurosciences, Medical Faculty, University of Bonn, Bonn, Germany; Molecular Physiology, Center for Integrative Physiology and Molecular Medicine (CIPMM), University of Saarland, Homburg, Germany; German Center for Neurodegenerative Diseases (DZNE), Bonn, Germany; Institute of Neurology, University College London, London, United Kingdom; Institute of Molecular Psychiatry, Medical Faculty, University of Bonn, Bonn, Germany

**Keywords:** NG2 glia, Kir4.1, hippocampus, myelination LTP, object recognition

## Abstract

NG2 glia represent a distinct type of macroglial cells in the CNS and are unique among glia because they receive synaptic input from neurons. They are abundantly present in white and grey matter. While the majority of white matter NG2 glia differentiates into oligodendrocytes, the physiological impact of grey matter NG2 glia and their synaptic input are ill defined yet. Here we asked whether dysfunctional NG2 glia affect neuronal signaling and behavior. We generated mice with inducible deletion of the K^+^ channel Kir4.1 in NG2 glia and performed comparative electrophysiological, immunohistochemical, molecular and behavioral analyses. Focussing on the hippocampus, we found that loss of the Kir4.1 potentiated synaptic depolarizations of NG2 glia and enhanced the expression of myelin basic protein. Notably, while mice with targeted deletion of the K^+^ channel in NG2 glia showed impaired long term potentiation at CA3-CA1 synapses, they demonstrated improved spatial memory as revealed by testing new object location recognition. Our data demonstrate that proper NG2 glia function is critical for normal brain function and behavior.

## Introduction

NG2 glial cells, which have also been termed complex cells, GluR cells, synantocytes, oligodendrocyte precursor cells, or polydendrocytes (reviewed by ***Bergles et al***., 2010), represent a distinct type of macroglial cells in the CNS (***Peters***, 2004; ***Nishiyama et al***., 2009). They are unique among glia because they receive direct synaptic input from glutamatergic and GABAergic neurons (***Bergles et al***., 2000; ***Lin and Bergles***, 2004; ***Jabs et al***., 2005). NG2 glia are abundantly present in white and grey matter, make up about 5-10% of total glial cell numbers and display variability with regard to their functional properties and antigen profiles (***Trotter et al***., 2010; ***Degen et al***., 2012; ***Seifert and Steinhäuser***, 2018; ***Marisca et al***., 2020). They have a high proliferative potential and, in grey matter, many cells keep their NG2 phenotype throughout adulthood (***Dimou et al***., 2008; ***Kang et al***., 2010; ***Moshrefi-Ravasdjani et al***., 2017). Despite intense research, the physiological impact of grey matter NG2 glia and their synaptic innervation are yet largely unknown. This deficit is mainly due to the fact that NG2 glia express a similar set of ion channels and receptors as neurons, although often at a much lower density, which makes selective manipulation of the glial cells difficult. To circumvent this problem, cell-type specific genetic deletion strategies have been developed. Thus, it has been demonstrated that ablation of NG2 cells entails depressive-like behavior (***Birey et al***., 2015) and impairs microglial function (***Liu and Aguzzi***, 2020), and deletion of AMPA receptors in NG2 glia affected white matter myelination (***Kougioumtzidou et al***., 2017).

The neurophysiological fingerprint of NG2 glia is mainly determined by the inward rectifier K^+^ channel Kir4.1 that is developmentally upregulated and dominates the membrane conductance in the physiologically relevant voltage range (***Schröder et al***., 2002; ***Djukic et al***., 2007; ***Tang et al***., 2009; reviewed by ***Seifert and Steinhäuser***, 2018), which allows them to sense [K^+^]_o_ (***Maldonado et al***., 2013). Thus, one promising approach to learn about NG2 glia function is to delete this important signaling molecule, to disturb normal NG2 glia function and test whether this entails altered neuronal function. Recently, Larson et al. generated mice with tamoxifen-induced deletion of Kir4.1 in PDGFRα^+^ NG2 glia. As expected, Kir4.1-lacking NG2 glia were significantly depolarized, but their survival, proliferation and differentiation were not affected (***Larson et al***., 2018). A similar approach was used by Song et al. to show that deletion of Kir4.1 from NG2 glia contributes to myelin loss in a transient ischemic mouse model (***Song et al***., 2018).

These findings imply that dysfunctional NG2 glia can affect neuronal signaling and mouse behavior. We tested this hypothesis by using NG2 CreERT2 knockin mice to inducibly delete Kir4.1 from NG2 glia. Focussing on the hippocampus, we demonstrate that loss of the K^+^ conductance significantly potentiated synaptic depolarizations of the glial cells, which was accompanied by an upregulation of myelin basic protein (MBP) on the transcript and protein levels. Notably, while mice with targeted deletion of Kir4.1 in NG2 glia showed impaired LTP at CA3-CA1 synapses, they demonstrated improved spatial memory as revealed by testing object location recognition. Thus, our data demonstrate that proper NG2 glia function is required for normal brain function and behavior.

## Results

### Efficiency of Kir4.1 deletion

Experiments were performed with NG2ki-EYFP (***Karram et al***., 2008) and Kir4.1 fl/fl;NG2-CreERT2 x Rosa26-EYFP mice (hereinafter referred to as Kir4.1 ko mice). To estimate the efficiency of homologous recombination we used several approaches at different time points after tamoxifen injection. First, immunostaining for the reporter EYFP (***Srinivas et al***., 2001) and PDGFRα, a specific marker of NG2 glia (***Rivers et al***., 2008) was performed (Suppl. Fig. 1A). The number of PDGFRα^+^EYFP^+^ cells was determined and divided by the total number of PDGFRα^+^ cells. EYFP^+^ cells contacting blood vessels were excluded, because they are pericytes and lack PDGFRα (***Karram et al***., 2008; ***Huang et al***., 2014). Between 4 and 8 weeks after tamoxifen injection, 60-80% of PDGFRα^+^ cells in the hippocampus also expressed EYFP, both in controls (i.e. mice lacking floxed Kir4.1) and Kir4.1 knockout (ko) mice (3 mice each; Suppl. Fig. 1A, B). This proportion might have been underestimated because of the limited fluorescence intensity of recombined cells in Rosa26-EYFP reporter mice (***Huang et al***., 2014). Four weeks after tamoxifen treatment, the density of PDGFRα^+^ cells was similar between control and Kir4.1 ko mice within each of the hippocampal subregions analyzed (Suppl. Fig. 1A, C; Suppl. Table 1). In control mice, however, the density of NG2 glia decreased by about 40% in all hippocampal subregions 8 weeks after tamoxifen application. In Kir4.1 ko mice only the density of PDGFRα^+^ cells in the stratum moleculare of the DG was significantly reduced (Suppl. Fig. 1C; Suppl. Table 1). These developmental changes are in line with previous data and might be attributed to a limited differentiation of NG2 glia and the increase of brain volume (***Zhang et al***., 2005; ***Moshrefi-Ravasdjani et al***., 2017)

Second, an indirect measure of recombination efficiency was obtained through patch clamp recording 3-4 weeks after tamoxifen injection (Fig. 1). EYFP^+^ (i.e. recombined) cells were selected in the CA1 stratum radiatum of control (n = 82 cells, N = 9 mice) and Kir4.1 ko mice (n = 183, N = 17) and tested for the presence of Kir currents. Unexpectedly, in Kir4.1 ko mice two populations of fluorescent cells co-existed in the same slices, either still expressing (termed Kir4.1 wt cells) or lacking Kir currents (membrane conductance at -130 mV, <6 pA/mV; termed Kir4.1 ko cells) (Fig. 1A, B). Compared to EYFP^+^ cells from control mice (n = 82, N = 9) and Kir4.1 wt cells (n = 55, N = 17), which had resting potentials close to the K^+^ equilibrium potential and a membrane resistance of 50.8 to 81.2 MΩ, Kir4.1 ko cells were dramatically altered in their passive membrane properties. These cells lacking Kir4.1 were characterized by depolarized resting potentials (-61 mV, n = 128; N = 17) and a drastically increased input resistance reaching values of >2 GΩ (n = 118; N = 17), while membrane capacitance was unaffected (Fig. 1C-E).

**Figure 1.**
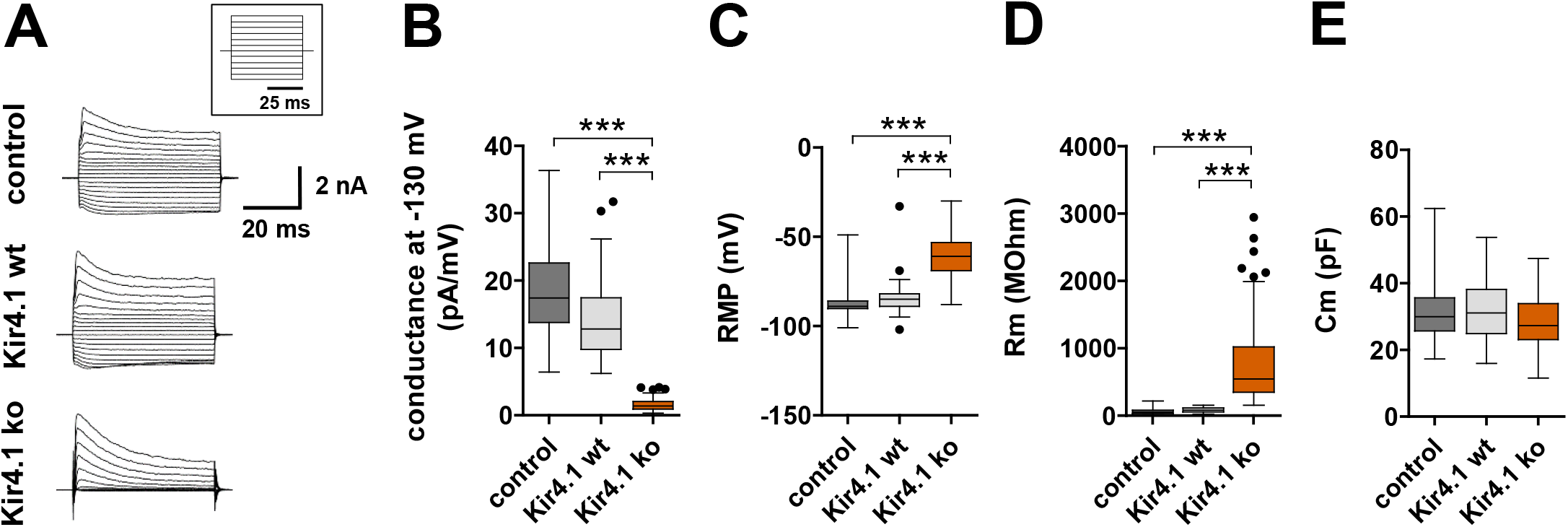
Kir4.1 channels determine passive membrane properties of NG2 glial cells. A) Examples of whole-cell current pattern of recombined NG2 glia cells in control and Kir4.1 ko mice evoked by de- and hyperpolarizing steps between -160 mV and +20 mV lasting for 50 ms. B) Cells with a conductances < 6 pA/mV at -130 mV were identified as Kir4.1 knockout (ko) cells (median Kir4.1 ko cells: 1.4 pA/mV, n = 128, N = 17). NG2 glia of Kir4.1 ko mice that still expressed Kir4.1 currents were termed Kir4.1 wildtype (wt) cells (median: 12.8 pA/mV, n = 55, N = 15). The membrane conductance of Kir4.1 wt cells resembled that of NG2 glia of control mice (median 17.4 pA/mV, n = 82, N = 9). C) In the absence of Kir4.1 the membrane potential of NG2 glia was depolarized to -61 mV compared to Kir4.1 wt (-85 mV) and control cells (-89 mV). D) The membrane resistance in cells lacking Kir4.1 (546 MΩ) was much higher than in Kir4.1 wt (81.2 MΩ) and control cells (50.8 MΩ). E) The membrane capacitance was not affected by deletion of Kir4.1 (median Kir4.1 ko cells, 27.3 pF; Kir4.1 wt cells, 31.1 pF; control cells, 30.0 pF). Kruskal-Wallis ANOVA with Dunńs test. Asterisks indicate statistically significant differences (*** p < 0.001).

In a third approach to estimate recombination efficiency, we determined Kir4.1 mRNA in FAC sorted NG2 glia from Kir4.1 ko and control mice. The quality of fluorescence activivated cell (FAC) sorting was evaluated in parallel experiments using wild type (C57BL6) mice as negative control and NG2ki-EYFP mice (***Karram et al***., 2008). With the latter, we defined a sorting window by the peak of EYFP fluorescence intensity at 527 nm as depicted in the sideward scatter of the FAC sorter. The fluorescence intensity of EYFP^+^ cells was at least 10 times higher than background fluorescence (Suppl. Fig. 2A). FAC sorted NG2 cells from hippocampi of NG2ki-EYFP mice (N = 10) were tested for cell type-specific mRNA expression (neurons, Rbfox3; microglia, Aif1; astrocytes, Aldh1L1; NG2 glia, PDGFRα). The housekeeping gene β-actin served as an internal standard. Gene expression ratios towards β-actin were determined and normalized to PDGFRα mRNA (N = 10). In FAC sorted hippocampal NG2 glia we found less than 2% of mRNAs coding for other cell types (Rbfox3, 0.79 ± 0.52%; Aif1, 1.11 ± 0.71%; AldH1L1, 0.12 ± 0.07%), confirming the high specificity of our FACS approach (Suppl. Fig. 2B). Kir4.1 mRNA was determined in recombined cells from the hippocampus of Kir4.1 ko and control mice, 4 weeks after tamoxifen. Threshold cycle differences, ΔCT, between Kir4.1 and β-actin were determined in individual samples and the gene ratio Kir4.1/β-actin was calculated (equation 1). In recombined NG2 cells of control mice the gene ratio was 0.102 ± 0.012 (N= 11), while in Kir4.1 ko mice, the ratio was 0.0136 ± 0.0049 (N = 8), i.e. 87% lower than in the controls (Suppl. Fig. 2C). The Kir4.1 mRNA amount in hippocampal NG2 cells from the NG2ki-EYFP mouse line was similar to recombined control cells in the hippocampus of NGCEcreERT2 mice (not shown).

To confirm the presence or absence of Kir4.1 mRNA in recombined NG2 glial cells with or without Kir currents, subsequent to patch clamp recording we harvested the cytoplasm of single fluorescent cells from the CA1 region of Kir4.1 ko mice. In cells without inward currents (Kir4.1 ko), Kir4.1 transcript was always absent (n = 15) while cells with inward currents (Kir4.1 wt) still expressed Kir4.1 mRNA (7/8 cells; Suppl. Fig. 2D). To test for functional expression of Kir4.1, Ba^2+^ (100 µM) was applied (focal pressure application). In control cells, membrane (chord) conductance decreased by almost 90% in Ba^2+^ solution (from 14.03 ± 4.13 to 1.78 ± 0.82 pA/mV, -130 mV, p = 0.02; n = 11) (Suppl. Fig. 2E), and similar Ba^2+^ sensitivity was found in Kir4.1 wt cells (reduction from 13.52 ± 5.26 to 1.61 ± 0.94 pA/mV, p = 0.02; n = 35) (Suppl. Fig. 2F). Notably, even in Kir4.1 ko cells the residual inward currents were Ba^2+^-sensitive (reduction from 1.46 ± 1.03 to 1.00 ± 0.52 pA/mV in Ba^2+^; p < 0.001; n = 49) (Suppl. Fig. 2G), which might suggest that part of the inward currents in NG2 glia were mediated by channels other than Kir4.1.

Together, these experiments revealed that in the hippocampus of Kir4.1 ko mice, tamoxifen-induced recombination led to the deletion of Kir4.1 channels in a majority of NG2 glial cells.

### Increased excitability of Kir4.1 deficient NG2 glial cells

Immature dentate granule cells (DGCs) express Kir channels at a lower density than mature DGCs, resulting in a higher input resistance and increased excitability upon synaptic input of the former (***Mongiat et al***., 2009). To test if Kir4.1 similarly determines input resistance and excitability of NG2 glial cells, miniature postsynaptic potentials (mPSPs) were recorded in the presence of TTX (0.5 µM). Indeed, mPSP amplitudes, rise and decay time constants in Kir4.1 ko cells (n = 27) significantly exceeded those of cells expressing Kir4.1 (Fig. 2A-D), which resembled changes seen when pharmacologically blocking Kir4.1 channels in NG2 glia (***Chan et al***., 2013). Upon quantal transmitter release, depolarizations of about 1.8 mV were detected in cells lacking Kir4.1 (n = 27), responses that were more than doubled compared to control (n = 13) and Kir4.1 wt cells (n = 22) (Fig. 2B). These depolarizations lasted several tens of ms in Kir4.1 ko cells while in cells still expressing Kir4.1, mPSPs declined to zero within 10 ms (Fig. 2A, D). Similarly, a slowdown of the rise time constant was observed in cells devoid of Kir4.1 (Fig. 2C). To estimate the frequency of mPSPs the inter event interval (IEI) was evaluated. Generally, mPSPs are rare events in NG2 glia (***Bergles et al***., 2000; ***Passlick et al***., 2016). In Kir4.1 ko cells, mPSPs occurred even less frequently, as indicated by an increased IEI compared to Kir4.1 wt cells although the difference to control cells was not significantly different (Fig. 2E). Together, these observations indicated that in NG2 glia, Kir4.1 critically determines membrane resistance and responsiveness to neuronal input.

**Figure 2.**
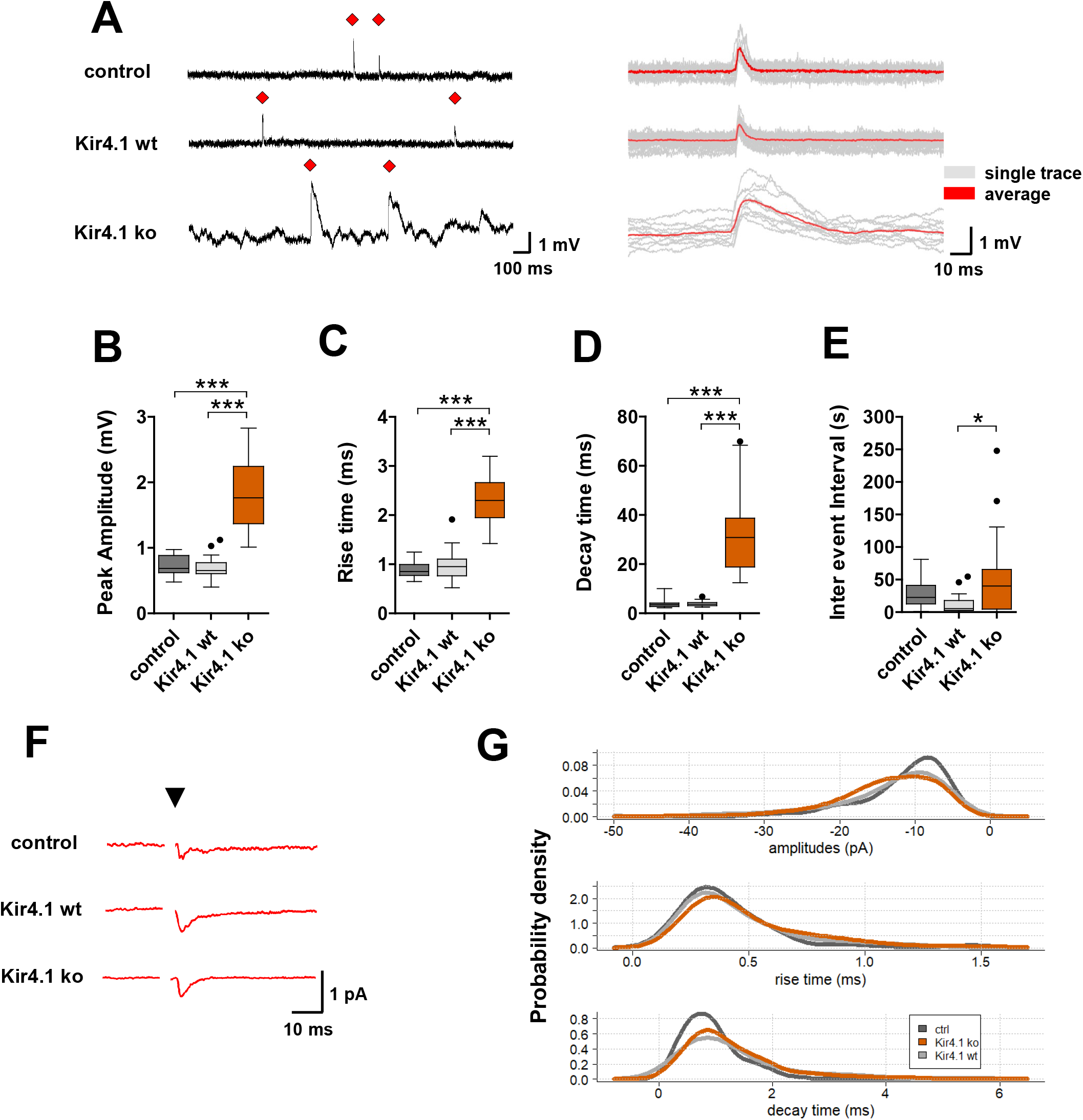
Increased amplitude and prolonged kinetics of mPSPs and ePSCs in NG2 glial cells lacking Kir4.1. A) Left: Example traces of mPSPs of control (top), Kir4.1 wt (middle) and Kir4.1 ko (bottom) NG2 glia cells. Red diamonds indicate individual events. Right: Corresponding superimposed single (grey) and average mPSP traces (red; average of 10-29 events). B) In Kir4.1 ko cells, peak mPSPs (n = 27, N = 15) were more than twice as high as in control (n = 13, N = 7) and Kir4.1 wt cells (n = 22, N = 12). C, D). Rise and decay time of mPSPs in Kir4.1 ko cells were significantly prolonged. E) The inter event interval of mPSPS was decreased in Kir4.1 ko cells compared to Kir4.1 wt cells. F) Averaged time course of NG2 glia eEPSCs upon stimulation of Schaffer collaterals. Stimulation pulse is indicated by black triangle. G) Probability density functions for amplitudes, rise and decay times of ePCSs in the presence of picrotoxin (150 µM). Number of control and Kir4.1 ko mice, N = 5 and 8. Asterisks indicate statistically significant differences (*** p < 0.001; * p = 0.014). See Tab. 2 for further details.

**Table 2.** Properties of miniature (A) and evoked PSPs (B) of control, Kir4.1 wt and Kir4.1 ko cells. Data is given as median and interquartile range (quartile 25% – quartile 75%), number of measures in square brackets (B).

| A | ACSF |  |  |
| --- | --- | --- | --- |
|  | Control | Kir4.1 wt | Kir4.1 ko |
| Amplitude (mV) | 0.69<br>(0.62 – 0.88) | 0.65<br>(0.61 – 0.77) | 1.76<br>(1.37 – 2.24) |
|  | Kir4.1 ko | < 2E-16 | - |
|  | control | 0.96 | < 2E-16 |
|  | ANOVA: |  | p < 2E-16 |
| Rise time (ms) | 0.85<br>(0.77 – 0.99) | 0.95<br>(0.76 – 1.11) | 2.30<br>(1.94 – 2.66) |
|  | Kir4.1 ko | 2E-09 | - |
|  | control | 0.7 | 3E-08 |
|  | Kruskal Wallis: |  | p = 3E-10 |
| Decay time (ms) | 3.49<br>(2.80 – 4.18) | 3.68<br>(3.11 – 4.40) | 30.84<br>(18.80 – 38.68) |
|  | Kir4.1 ko | 7E-09 | - |
|  | control | 0.86 | 1E-08 |
|  | Kruskal Wallis: |  | p = 2E-10 |
| Inter event interval (s) | 22.71<br>(12.64 – 41.05) | 5.27<br>(2.74 – 17.86) | 40.03<br>(4.68 – 65.71) |
|  | Kir4.1 ko | 0.01 | - |
|  | control | 0.06 | 0.87 |
|  | Kruskal Wallis: |  | p = 0.01 |
| Cells | 13 | 22 | 27 |

**Table 2. Properties of miniature (A) and evoked PSPs (B) of control, Kir4.1 wt and Kir4.1 ko cells.**
| B | ACSF |  |  | + PicTx |  |  |
| --- | --- | --- | --- | --- | --- | --- |
|  | Control | Kir4.1 wt | Kir4.1 ko | Control | Kir4.1 wt | Kir4.1 ko |
| Amplitude (pA) | 11.2<br>(6.8 – 16.3)<br>[68] | 18.0<br>(11.3 -25.1)<br>[212] | 18.1<br>(12.0 – 25.9)<br>[491] | 9.5<br>(7.5 – 13.2)<br>[87] | 11.1<br>(8.2 – 16.5)<br>[171] | 12.8<br>(8.8 – 16.7)<br>[227] |
|  | Kir4.1 ko | 0.23 | - | Kir4.1 ko | 0.17 | - |
|  | control | 2E-06 | 5E-09 | control | 0.12 | 6E-03 |
|  | Kruskal Wallis: |  | p = 1E-08 | Kruskal Wallis: |  | p = 8E-03 |
| Rise time (ms) | 0.34<br>(0.27 – 0.41)<br>[54] | 0.42<br>(0.34 – 0.54)<br>[173] | 0.47<br>(0.36 – 0.61)<br>[362] | 0.34<br>(0.28 – 0.49)<br>[80] | 0.37<br>(0.28 – 0.55)<br>[154] | 0.41<br>(0.31 – 0.61)<br>[196] |
|  | Kir4.1 ko | 6E-04 | - | Kir4.1 ko | 0.09 | - |
|  | control | 2E-02 | 2E-07 | control | 0.31 | 0.02 |
|  | Kruskal Wallis: |  | p = 2E-07 | Kruskal Wallis: |  | p = 0.01 |
| Decay time (ms) | 0.71<br>(0.43 – 1.39)<br>[59] | 1.66<br>(0.92 – 2.82)<br>[181] | 1.62<br>(1.09 – 2.48)<br>[423] | 0.86<br>(0.56 – 1.07)<br>[78] | 1.04<br>(0.64 – 1.57)<br>[159] | 1.02<br>(0.73 – 1.61)<br>[213] |
|  | Kir4.1 ko | 7E-09 | - | Kir4.1 ko | 0.51 | - |
|  | control | 0.79 | 2E-10 | control | 7E-03 | 1E-03 |
|  | Kruskal Wallis: |  | p = 3E-10 | Kruskal Wallis: |  | p = 1E-03 |
| Cells | 4 | 12 | 6 | 5 | 7 | 8 |

Both glutamatergic and GABAergic input depolarizes NG2 glia (***Lin and Bergles***, 2004; ***Jabs et al***., 2005; ***Passlick et al***., 2013). To isolate glutamatergic from GABAergic input, the bath solution was supplemented with the GABA_A_R blocker picrotoxin (150 µM). Peak amplitude and kinetics (rise time and decay time) of mPSPs were not affected by the blocker (Suppl. Fig. 3A-C) while the inter event interval in Kir4.1 wt cells was decreased in picrotoxin (Suppl. Fig. 3D) (Suppl. Table 2). Co-application of picrotoxin and NBQX (10 µM), a blocker of AMPA/kainate receptors, fully abolished mPSPs (not shown, control cells, n = 3; Kir4.1 wt, n = 5; Kir4.1 ko, n = 3).

Minimal stimulation of Schaffer collaterals was applied to test whether deletion of Kir4.1 in NG2 glia also affects evoked postsynaptic currents (ePSCs) (Fig. 2F). Compared to control cells, in Kir4.1 ko cells rise time, decay time and amplitudes of the ePSCs increased to 138, 228 and 161%, respectively (Table 2B). In the presence of picrotoxin (150 µM), similar changes were found (rise time, decay time and amplitudes, increase to 121, 119 and 134%, respectively; see Table 2B for details). The probability density functions of the eEPSC parameters were shifted towards higher excitability of Kir 4.1 ko cells (Fig. 2G). Taken together, both spontaneous mPSPs and ePSCs revealed significant increases in excitability of NG2 glial cells after functional knock out of Kir 4.1 channels.

### Impaired LTP in mice with Kir4.1 deficient NG2 glia

To test whether the increased responsiveness of NG2 glia in the hippocampus of Kir4.1 ko mice affects neuronal signaling, field excitatory postsynaptic potentials (fEPSPs) were recorded in the stratum radium of the CA1 region upon stimulation of the Schaffer collaterals (Fig. 3A). Basal excitability in the hippocampus was similar in control and Kir4.1 ko mice as judged by the unchanged input-output curves and by stimulus intensities required to evoke half-maximum fEPSP amplitudes (Suppl. Fig. 4A, B). Next we investigated long-term potentiation (LTP) of transmission at Schaffer collateral - CA1 pyramidal cell synapses. A standard theta burst protocol was used to induce LTP in slices of control (9 slices, N = 3) and Kir4.1 ko mice (30 slices, N = 12). We noted reduced LTP (by 17.6% at 25-30 min after TBS) in slices obtained from Kir4.1 ko mice (Fig. 3B, C), indicating that NG2 glial cells influence neuronal plasticity. This effect was not caused by changes in presynaptic release probability because the paired pulse ratios before and after LTP induction were similar in control (9 slices, N = 3) and Kir4.1 ko mice (24 slices, N = 9) (Fig. 3D, E).

**Figure 3.**
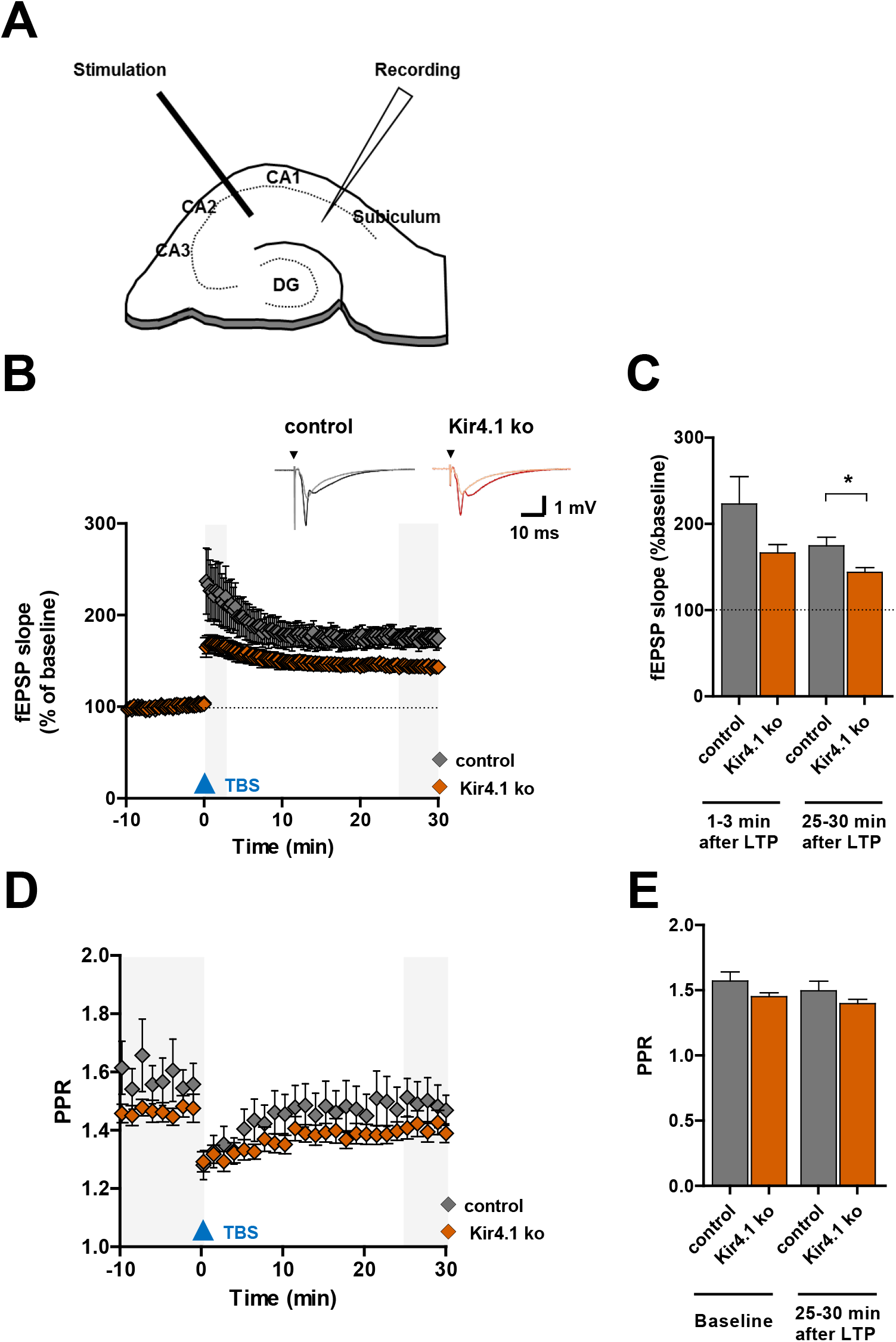
TBS-induced LTP is impaired in mice with NG2 glia-specific Kir4.1 deficiency. A) Position of stimulation and recording electrodes in hippocampal slices during field potential recording. B) Time course of normalized fEPSP slopes of control (n = 9 slices, N = and Kir4.1 ko mice (n = 30, N = 12) before and after induction of LTP by 3 trains of theta-burst stimulation (TBS, blue triangle). Insets show representative fEPSPs from control and Kir4.1 ko mice (before TBS, grey and orange; 25-30 min after TBS, black and red). Stimulation pulse is indicated by black triangles. C) LTP was impaired in hippocampal slices from Kir4.1 ko mice, 25-30 min after TBS (143.98 ± 5.37% vs 174.71 ± 9.84%, Mann-Whitney U test, p = 0.014). D) Time course of the paired pulse ratio (PPR) of control (n = 9, N = 3) and Kir4.1 ko mice (n = 24, N = 9). E) PPR was affected by Kirr4.1 ko mice, neither before (1.57 ± 0.07 vs 1.45 ± 0.03, two-sample t-test) nor 25-30 min after TBS-induced LTP (1.50 ± 0.07 vs 1.40 ± 0.03, two-sample t-test). Mean ± SEM.

### Proliferation and differentiation of NG2 glia are only slightly affected after deletion of Kir4.1

To clarify whether inducible deletion of Kir4.1 in NG2 glial cells affects their proliferation or differentiation, immunohistochemical analysis of the proliferation marker Ki67 and a marker for mature oligodendrocytes, GSTpi, was combined with molecular analysis. Recombined EYFP^+^ NG2 glia were identified with an GFP antibody. In different layers of the hippocampal CA1 region and the molecular layer of the dentate gyrus (DG), only few Ki67^+^EYFP^+^ cells were found 8 weeks post injection (wpi; <10% of all recombined cells), and there was no difference between control and Kir4.1 ko mice (Fig. 4A, B; Suppl. Table 3). In control animals, however, the total density of Ki67^+^ cells in the stratum lacunosum-moleculare was more than twice as high as in the radiatum, and deletion of Kir4.1 in NG2 glia induced an increase in total Ki67^+^ cell density in the stratum radiatum (Fig. 4C, Suppl. Table 3).

**Figure 4.**
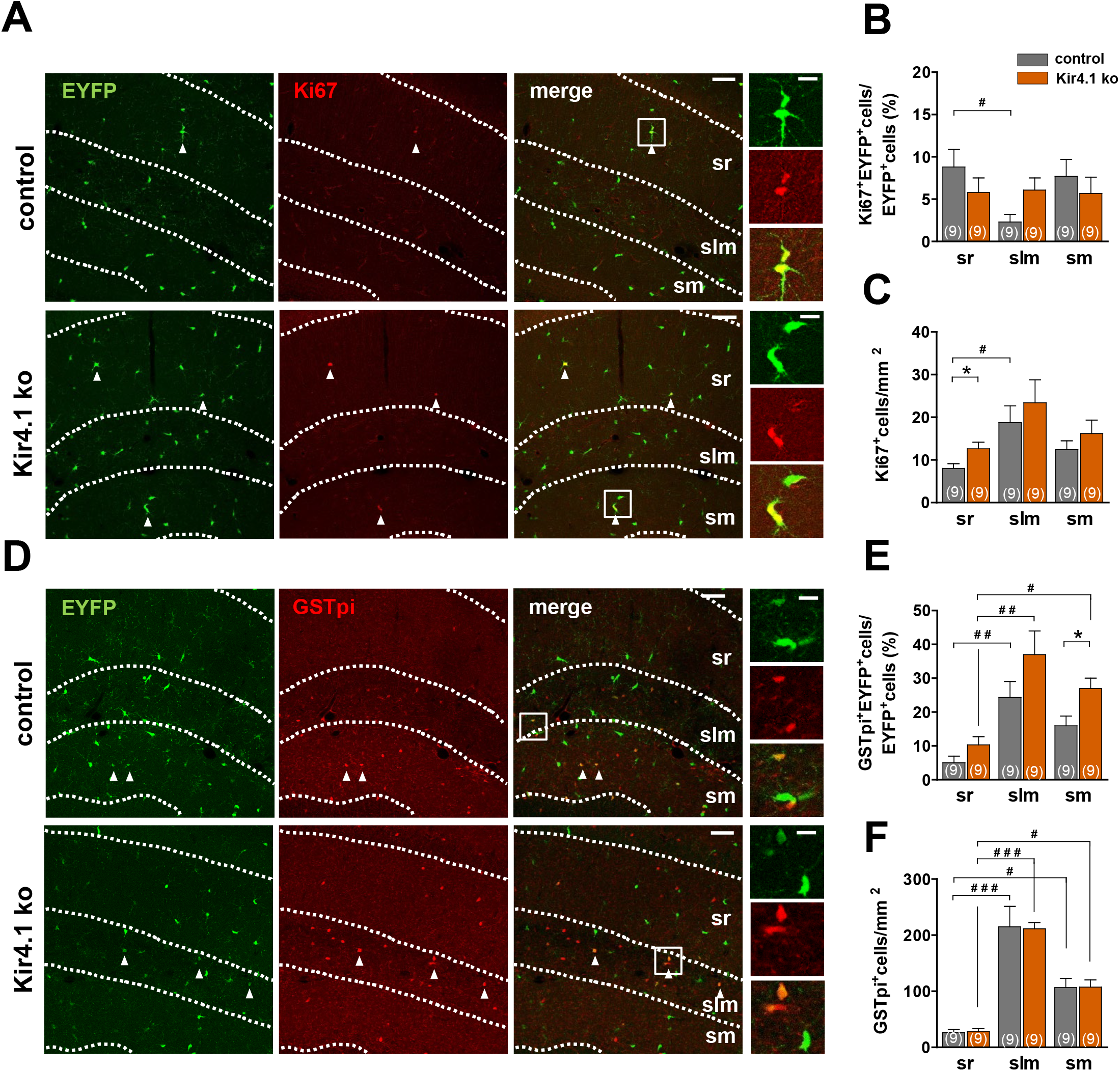
Proliferation and differentiation of NG2 glia is not affected in Kir4.1 ko mice. A) *Hippocampal* sections from control and Kir4.1 ko mice were immunostained with EYFP (green) and Ki67 (red). EYFP^+^Ki67^+^ cells are indicated by arrowheads. Boxed areas are given at higher resolution to the right. Scale bar 60 and 15 µm (insets). B) The proportion of Ki67^+^EYFP^+^ cells among all EYFP^+^ cells was similar in control and Kir4.1 ko mice. In control mice, the rate of NG2 glia proliferation in the sr was higher than in the slm (p = 0.037). C) In the sr of Kir4.1 ko mice the density of Ki67^+^ cells was higher than in the controls (12.60 ± 1.47 vs. 8.0 ± 1.04, p = 0.038). Within the control group, the density of proliferating cells in the slm was higher than in the sr (p = 0.038). D) Sections were immunostained with EYFP (green) and GSTpi (red). EYFP^+^GSTpi^+^ cells are indicated by arrowheads. Boxed areas are shown at higher resolution to the right. Scale bars 60 and 15 µm (insets). E). In Kir4.1 ko mice, the proportion of recombined NG2 glia (EYFP^+^) differentiating into oligodendrocytes (EYFP^+^ GSTpi^+^ cells) in the sr was lower than in the sm of the DG ( p = 0.023) and slm (p = 0.003). Layer-specific differences were also found in control mice (sr vs slm, p = 0.004). In the sm, deletion of Kir4.1 in NG2 glia stimulated differentiation of these cells into oligodendrocytes (p = 0.018). F) The density of GSTpi^+^ cells was not different between control and Kir4.1 ko mice. However in both genotypes higher values were observed in the slm compared to the sr (control, p < 0.001; Kir4.1 ko, p < 0.001) and between sr and sm (control, p = 0.018; Kir4.1 ko, p = 0.041). Significant differences between control and Kir4.1 ko mice are indicated by asterisks (* p < 0.05; Mann-Whitney U Test), differences between layers within a genotype by hash marks (# p < 0.05; ## p < 0.01; ### p < 0.001; Kruskal-Wallis ANOVA with Dunńs test). Mean ± SEM. Number of mice is given in bar graphs. For further details see Suppl. Table 3.

Next, immunostaining against gluthathione S-transferase pi (GSTpi), an enzyme expressed by oligodendrocytes (***Tansey et al***., 1991) was performed to test for enhanced NG2 glia differentiation after targeted Kir4.1 deletion. Control stainings using NG2ki-EYFP mice (***Karram et al***., 2008) confirmed the absence of GSTpi expression in EYFP^+^ NG2 glia of the hippocampus (not shown). Eight wpi, 5 - 37% of all recombined NG2 glial cells in the hippocampus of control and Kir4.1 ko mice were positive for GSTpi. Targeted deletion of Kir4.1 induced an increase in GSTpi^+^EYFP^+^ cells in the stratum moleculare of the DG while the differences in the other layers were not statistically significant (Fig. 4D, E; Suppl. Table 3). The density of GSTpi^+^ cells was not affected in Kir4.1 ko mice (Fig. 4F). Thus, postnatal deletion of Kir4.1 in NG2 glia provoked no or only subtle changes in the proliferation or differentiation of those cells.

### Increase of myelin-specific gene transcripts and protein in Kir4.1 ko mice

Although gray matter NG2 glial cells do not form myelin or express myelin-related protein (***Diers-Fenger et al***., 2001), they do express corresponding transcripts, e.g. for myelin-associated glycoprotein (MAG), myelin basic protein (MBP) or 2,3-cyclic nucleotide 3-phosphodiesterase (CNP) (***Ye et al***., 2003; ***Moshrefi-Ravasdjani et al***., 2017). In this study, we wondered if targeted deletion of Kir4.1 in NG2 glia affects expression of MAG and MBP mRNA in FAC sorted cells. Three wpi, the MAG-to-β-actin expression ratio in hippocampi of control and Kir4.1 ko mice was 1.15 ± 0.17 (N = 12) and 1.69 ± 0.11 (N = 8). Similarly, the ratio of MBP-to-β-actin mRNA expression in control mice (2.32 ± 0.42, N = 12) was lower than after Kir4.1 deletion (5.04 ± 0.44, N = 8) (Fig. 5A).

**Figure 5.**
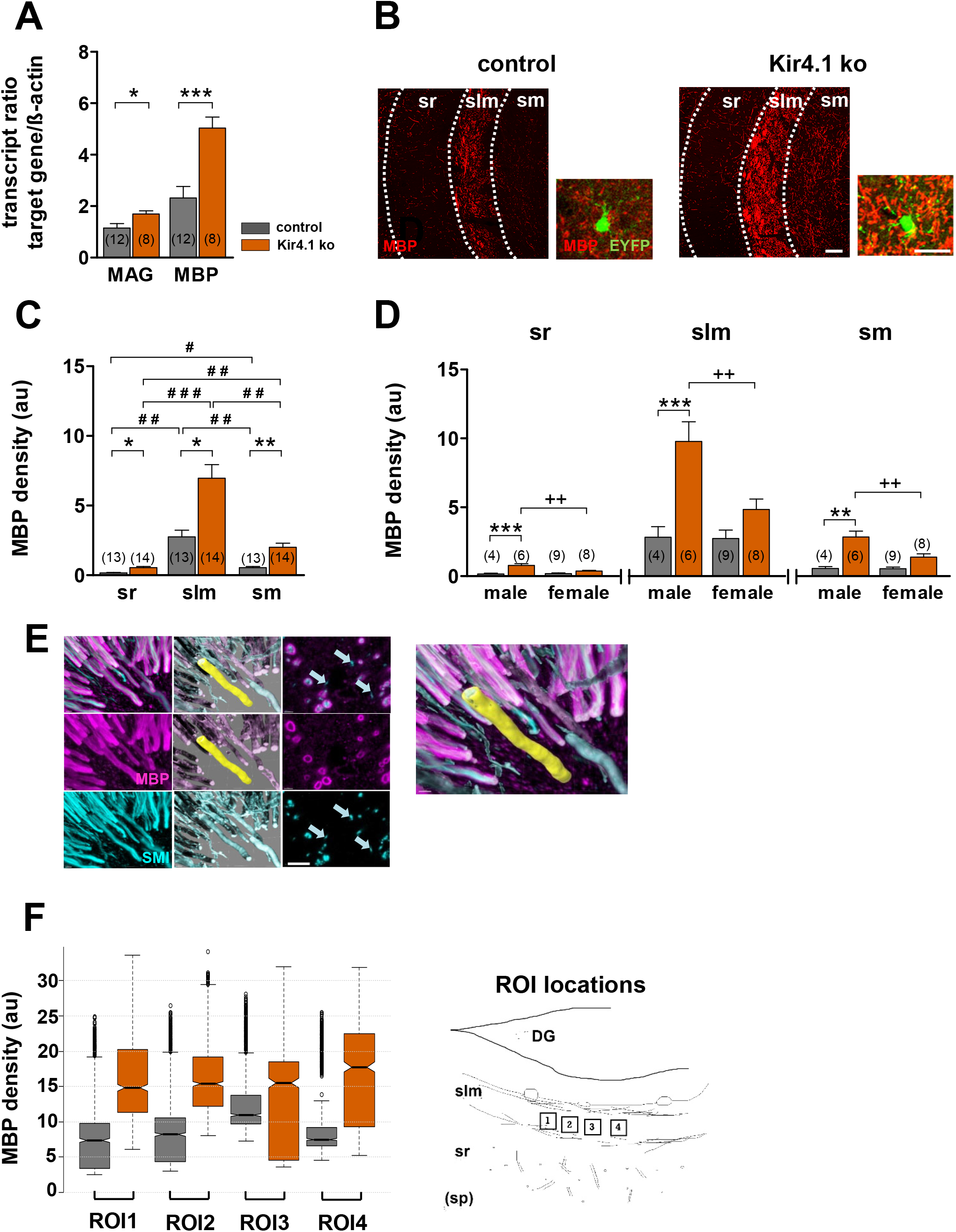
Enhanced MBP expression in mice with NG2 glia-specific deletion of Kir4.1. A) Transcript analysis of FACsorted recombined NG2 glia of Kir4.1 ko mice revealed a significant upregulation of MAG (* p < 0.05; t-test) and MBP (*** p < 0.001; t-test). B) Hippocampal sections from control and Kir4.1 ko mice were immunostained for MBP (red). Blowups show recombined NG2 glial cells (EYFP^+^) expressing MBP. Scale bars, 50 and 15 µm (blowups). C) MBP immunoreactivity was determined as fluorescent intensity per area (density) in slices from control (n=34) and Kir4.1 ko mice (n=41). Data were normalized to the respective median of MBP density. Location and genotype were significantly different for the parameter MBP density (2-way 3x2 ANOVA, location p < 0.05, genotype p < 0.05). In both, Kir4.1 ko and control mice, maximal values were observed in the slm. Significant differences between control and Kir4.1 ko mice within layers are indicated by asterisks, differences between layers within one genotype by hash tags (2-way 3x2 ANOVA followed by paired comparisons with Bonferroni adjustment) (*, # p < 0.05; **, ## p < 0.01; ### p < 0.001). D) MBP immunoreactivity in male Kir4.1 ko mice was not only stronger than in male control mice (** p < 0.01; *** p < 0.001) but also compared to female Kir4.1 ko mice in all three layers (^++^ p < 0.01) (2-way 2x2 ANOVA with post hoc Tukey HSD for each layer). Mean ± SEM. Number of mice is given in bar graphs. E) Subcellular analysis of MBP and SMI expression revealed by expansion microscopy (control mouse). 3D representations of immunolabeling against myelin sheaths (MBP, purple), axons (SMI, turquoise) and computer-engineered matched volumes constructed by isosurfaces (semitransparent layers of the same color). The yellow structure is one exemplary highlighted MBP volume. Note the shape of a hollow cylinder. Shown are 3D visualizations (cavalier projections, 14.5 x 10.3 x 11 µm^3^) of immunofluorescence (left column) and fitted isosurfaces (middle). The right column displays one single xy-plane taken from the corresponding confocal picture stack of immunostainings. The top row depicts the overlay of MBP and SMI structures, middle row MBP staining and the bottom row gives the SMI staining pattern. Arrows indicate examples of unmyelinated nerve fibers. Scale bars, 1.15 µm. Images were taken from a control mouse. Right: Enlarged detail of superposed immunostainings and corresponding isosurface-based fitted particles. F) Statistical analysis of MBP data obtained with expansion microscopy. Depicted are notched box plots of the density (summed intensity divided by volume, left) of the MBP isosurface particles. Data were collected from a total of 4 slices (2 control mice, grey, and 2 Kir4.1 ko mice, orange). In each slice, 4 ROIs were analyzed (position of ROIs is shown on the right). A total of 5107 and 3828 particles were analyzed for control and Kir4.1 ko mice, respectively. In each individual ROI of slices from Kir4.1 ko mice, the density was significantly higher than in the corresponding controls (Kruskal Wallis test) (ROI1, increase to 201%, p < 0.001; ROI2, to 188%, p < 0.001; ROI3, to 141%, p = 0.028; ROI4 to 237%, p < 0.001). Data are give as median ± SEM. Box 25-75 percentile, whiskers ± 1.5 IQR, notches ± 1.58 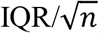, dots show outliers.

These gene expression data were further confirmed by immunostainings for MBP in Kir4.1 ko mice. Immunoreactivity levels were normalized to the stratum lacunosum moleculare of Kir4.1 ko mice (n = 41 slices; N = 14 mice). In line with the transcript data, 8 wpi strongly elevated MBP levels were observed in all layers of Kir4.1 ko mice as compared to control mice (n = 37; N = 13) (stratum radiatum, 0.542 ± 0.312 vs 0.173 ± 0.14; stratum lacunosum moleculare, 6.96 ± 3.68 vs 2.76 ± 1.68; statum moleculare 2.00 ± 1.11 vs 0.543 ± 0.329) (Fig. 5B, C). Notably, this genotype-dependent increase was confined to male mice (Kir4.1 ko, n = 17 slices; N = 6 mice; control mice, n = 12; N = 4) (stratum radiatum: Kir4.1 ko, 0.779 ± 0.321 vs. control, 0.162 ± 0.112, stratum lacunosum moleculare: Kir4.1 ko, 9.79 ± 3.49 vs. control, 2.83 ± 1.52; Kir4.1 ko statum moleculare, 2.84 ± 1.03 vs. control, 0.552 ± 0.289). Thus, male Kir4.1 ko mice showed a stronger increase in MBP immunoreactivity than females (n = 24; N = 8) in these hippocampal subregions (female Kir4.1 ko: stratum radiatum, 0.363 ± 0.152; stratum lacunosum moleculare, 4.84 ± 2.12; statum moleculare 1.38 ± 0.693) (Fig. 5D).

To get further insight as to how targeted deletion of Kir4.1 affects myelination, axons and myelin sheaths were stained with antibodies against SMI312 and MBP and analyzed at higher spatial resolution using expansion microscopy (***Chen et al***., 2015; ***Chozinski et al***., 2016). Analysis was confined to the stratum lacunosum moleculare because of the high MBP immunoreactivity seen in this layer (Fig. 5B-D). Based on the immunostainings, 3D isosurfaces of axons (SMI312) and myelin sheaths (MBP) were constructed, further referred to as particles (Fig. 5E). For MBP particles, we calculated volume, total fluorescence intensity and the density of fluorescence intensity. In Kir4.1 ko mice, in all individual ROIs placed into stratum lacunosum moleculare (Fig. 5F, right) a reduction in MBP volume (decrease of the averaged medians to 74 ± 36%, n = 8 ROIs from 2 slices, p = 0.02) was found while cumulative MBP fluorescence intensity did not change (p = 0.09) (not shown). Notably, the cumulative fluorescence density of MBP particles was almost doubled in all individual ROIs in Kir4.1 ko vs. control mice (increase of the averaged medians to 192 ± 34%, p < 0.001) (for averages per ROI, see Fig. 5F). Removing all data outliers (± 1.5 IQR) did not alter the effect size of fluorescence density (189 ± 34%, p < 0.001). In conclusion, these data indicate a significant increase in MBP density within the myelin sheaths in the hippocampus of Kir4.1 ko mice.

### Mice with Kir4.1 deficient NG2 glia show improved novel object location recognition

Several tests were performed to investigate whether dysfunctional NG2 glial cells affect mouse behavior. We assessed spatial working memory of the animals using the Y-maze test, where the ratio of alternations correlates with the strength of working memory (N = 17 and 19 for control and Kir4.1 ko mice, respectively). Neither genotype nor sex influenced the behavior of the mice in this paradigm, and no interaction between sex and genotype was found (Fig. 6A).

**Figure 6.**
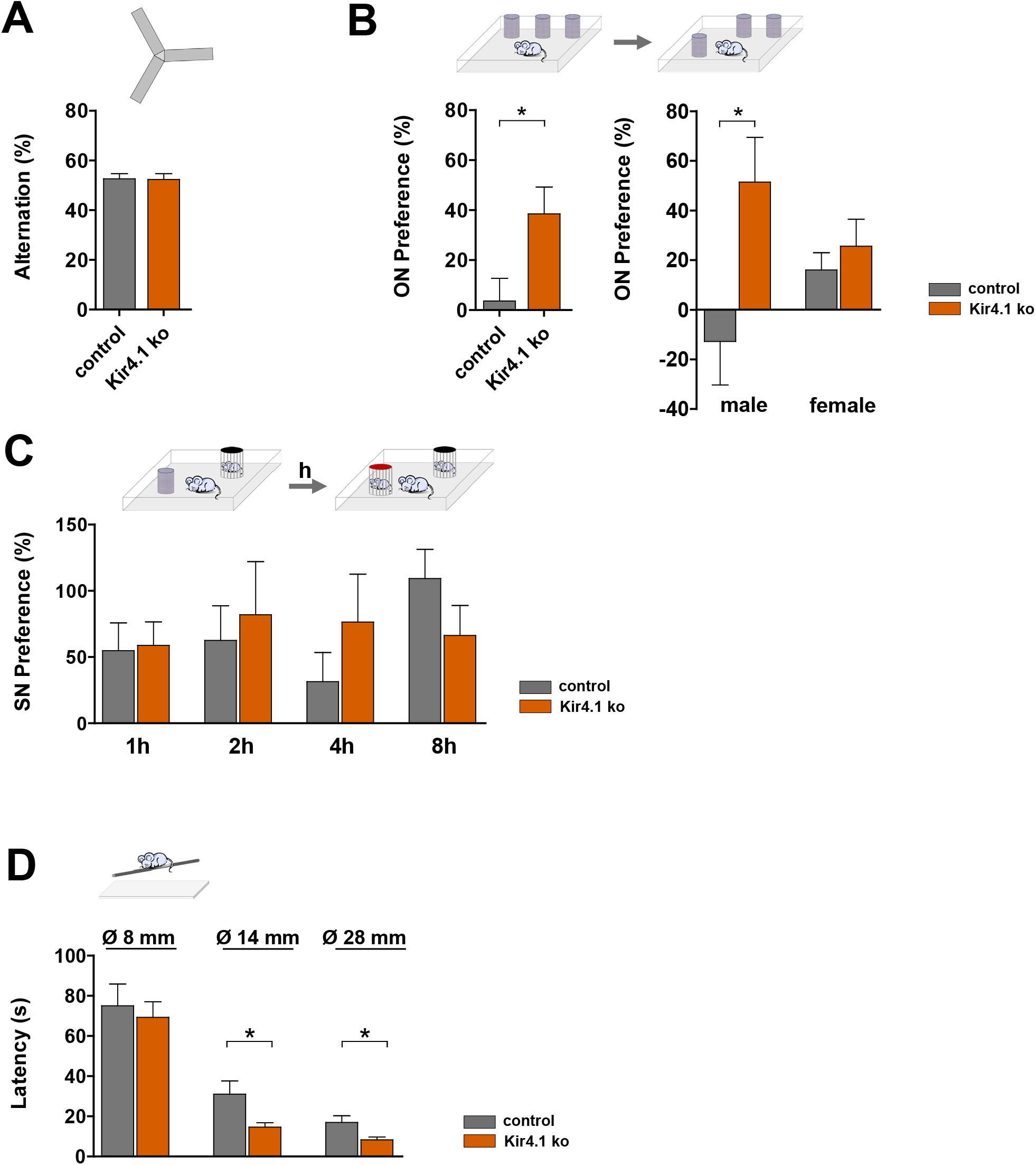
NG2 glia-specific deletion of Kir4.1 improves performance in new object location recognition and motor coordination tests. A) Working memory was tested in a Y-maze paradigm. Genetic deletion of Kir4.1 did not influence spatial working memory as the proportion of alternation was not significant between Kir4.1 ko and control mice (control, N = 17; Kir4.1 ko mice, N = 19). B) Novel object location recognition test. Object novelty (ON) preference, an indicator of spatial memory, was significantly higher in Kir4.1 ko mice (38.50 ± 10.34%, N = 14) than in control mice (3.66 ± 8.69%, N = 14) (left). Sex (p = 0.93) and genotype (p = 0.016) yielded no interaction between these two factors (p = 0.055) but the posthoc test indicated that the difference between genotypes was confined to male mice (p = 0.017; females, p = 0.954) (right). C) Partner recognition test. Novelty preference as an indicator of social memory was tested at different inter-trial times (1 h, 2 h, 4 h, 8 h). No difference was detected between Kir4.1 ko (N = 19) and control mice (N = 16). D) Beam walking test on rods with 28 mm, 14 mm and 8 mm diameter. The thinner the rod, the more difficult the balancing task is. Kir4.1 ko mice showed lower latencies than control mice on rods with a diameter of 28 mm (8.33 ± 1.38, N = 18 vs 16.94 ± 3.37, N = 17; p = 0.022) and 14 mm (14.63 ± 2.18, N = 19 vs 31.57 ± 6.57, N = 17; p = 0.017), suggesting better motor coordination. 2-way ANOVA with Tukey HSD. Mean ± SEM. Asterisks indicate statistically significant differences (* p < 0.05).

Testing novel object location recognition (NOLR) is a sensitive tool to evaluate spatial memory. Animals, which remember the original location of an object spend more time investigating the object in a new position, thus showing novelty preference. Notably, novelty preference values indicated a significantly improved recognition ability of Kir4.1 ko mice (N = 14 for control and Kir4.1 ko mice, p = 0.016) (Fig. 6B left), and this difference between genotypes was exclusively seen in male mice (p = 0.017) (Fig. 6B right).

To test social memory, we applied a partner recognition test. Similar to the novel object location recognition test, a positive novelty preference value is an indicator of recognition. Using 1 h inter-trial intervals we found a significant sex effect (p = 0.007), because females showed a higher partner recognition ability. However, the genotype of the animals did not influence partner recognition, and sex - genotype interaction was also not present. Using longer intervals (2 h, 4 h, 8 h) neither genotype nor sex influenced social memory (Fig. 6C). Sex - genotype interaction was also not present. Thus, our result suggests that mice with targeted deletion of Kir4.1 in NG2 glia have similar social memory as control mice.

Finally, motor and coordination abilities of male and female controls and mice with Kir4.1 deletion in NG2 glia were compared using the beam walking test. Here, a lower latency to reach the goal box indicates better coordination ability. On the wide (28 mm diameter) and middle-sized beams (14 mm), Kir4.1 ko mice reached the goal box significantly faster than the controls (14 mm, p = 0.03; 28 mm, p = 0.03) (Fig. 6D middle, right). Sex effects or genotype - sex interactions were not present. Testing the animals on the thinnest rod (8 mm diameter) we found a significant interaction between sex and genotype (p < 0.01) but no effect of genotype or sex alone.

Together, our data reveal that NG2 glia-specific deletion of Kir4.1 improves spatial memory and motor abilities. Interestingly, deletion of Kir4.1 did not lead to a generally improved learning and memory, because working memory or social declarative memory remained unchanged in the ko mice.

## Discussion

### Targeted deletion of Kir4.1 channels amplifies synaptic input onto NG2 glia

In contrast to astrocytes and oligodendrocytes, hippocampal NG2 glia are not coupled by gap junctions (***Wallraff et al***., 2004; ***Griemsmann et al***., 2015) and spatial buffering of K^+^, which utilizes Kir4.1 channels in the other glial cell types (***Wallraff et al***., 2006; ***Larson et al***., 2018; ***Schirmer et al***., 2018) is unlikely to occur. Nevertheless, a significant developmental upregulation of inward rectifying currents has been observed in NG2 glia (***Kressin et al***., 1995; ***Maldonado et al***., 2013), which is mostly mediated by Kir4.1 channels (***Djukic et al***., 2007; ***Tang et al***., 2009; ***Song et al***., 2018). K^+^ fluctuations induced by neuronal activity may be sensed by adult grey matter NG2 glia, and trains of neuronal stimulation may depolarize adult NG2 glia and modulate synaptic currents (***Maldonado et al***., 2013). Variable Kir current amplitudes were not only observed during development but also between brain regions, with white matter NG2 glia displaying lower current densities than cells in grey matter (***Chittajallu et al***., 2004). In our study, genetic deletion of Kir4.1 channels led to a tenfold increase in input resistance, similar to what was observed when Ba^2+^ was used to block Kir channels (***Chan et al***., 2013). The latter study has shown that increase in R_M_ in thin processes broadens the time window of temporal summation of synaptic signals in NG2 glia, as also predicted in a cable model. Indeed, we observed much larger and longer spontaneous and evoked PSPs in Kir4.1 deficient cells indicating enhanced summation of synaptic input onto NG2 glia, which might activate voltage-activated Ca^2+^ channels (***Haberlandt et al***., 2011; ***Sun et al***., 2016). Our study confirms that Kir4.1 channels mediate the main K^+^ conductance in adult NG2 glia. These channels not only determine passive membrane properties but also regulate the efficiency of NG2 glia depolarization upon synaptic input.

In the naive hippocampus, a single NG2 glial cell receives input from less than 20 glutamatergic neurons, and the GABAergic input is even weaker (***Bergles et al***., 2000; ***Haberlandt et al***., 2011; ***Sun and Dietrich***, 2013). The increased interevent intervals of mEPSPs after Kir4.1 deletion may indicate even lower NG2 glia synaptic connectivity, and loss of connectivity seems also to accompany postnatal developmental (***Passlick et al***., 2016). Notably, short-term synaptic plasticity, which is a measure of vesicle release probability and related to presynaptic Ca^2+^ levels (***Zucker and Regehr***, 2002), was not affected in Kir4.1 deficient NG2 glia synapses.

### Deletion of Kir4.1 in NG2 glia does not affect proliferation and differentiation but increases MBP density

Global knockout of Kir4.1 resulted in reduced body development, tremor and premature lethality. In the CNS, the development of oligodendrocytes was impaired and vacuolization of white matter tracts was observed, accompanied by reduced myelination and axonal degeneration (***Neusch et al***., 2001). Conditional, GFAP promoter-driven Kir4.1 ko mice developed ataxia, seizures, and the number of NG2 glia (at that time still called ’complex cells’, see ***Bergles et al***., 2010) in the hippocampus was profoundly reduced. NG2 glia remaining in those ko mice lost their negative resting potentials and displayed increased membrane resistance (***Djukic et al***., 2007). Notably, inducing ablation of Kir4.1 from NG2 glia (this study) or oligodendrocytes (***Larson et al***., 2018) only at postnatal week 3-4, i.e. after onset of myelination, overcame this burdened phenotype.

The amplified synaptic input after Kir4.1 deletion did not influence proliferation of NG2 glia, which agrees with a previous study using a BrdU assay in PDGFRα-CreER-based Kir4.1 ko mice (***Larson et al***., 2018). Interestingly, reduced numbers of BrdU^+^/Olig2^+^ cells were observed in the spinal chord of newborn mice with constitutive Olig2-Kir4.1 deletion, suggesting that lower proliferation entailed an earlier onset of myelination. The authors also reported reduced numbers of progenitor cells, but enhanced MBP transcript levels in newborn Kir4.1-deficient NG2 glia (***Schirmer et al***., 2018). Our NG2-induced deletion of Kir4.1 at the adult stage increased MBP not only on the transcript but also on the protein level. Tamoxifen-induced recombination in NG2-CreERT2 mice allowed us to track cell fate. We found some cells, both in control and Kir4.1 ko mice, co-expressing GSTpi with EYFP 8 weeks after tamoxifen injection, indicating differentiation of NG2 glia into oligodendrocytes (***Tamura et al***., 2007; ***Huang et al***., 2014). Differentiation differed between hippocampal layers, being particularly evident in the stratum lacunosum moleculare. Variable expression of growth factors, such as PDGF, FGF-2 NT-3 and NGF, or its receptors, might account for the subregional differences in proliferation and differentiation (***Barres et al***., 1994; ***Cohen et al***., 1996; ***Mason and Goldman***, 2002; ***Wilson et al***., 2003).

However since neither the proliferation rate of NG2 glia nor the density of GSTpi^+^ cells differed between genotypes we conclude that targeted deletion of Kir4.1 did not stimulate NG2 glia differentiation in the hippocampus. This is in line with a study by Larsen et al. (2018) who found no changes in NG2 glia differentiation 2 weeks after tamoxifen-induced Kir4.1 from PDGFRα-expressing cells. Thus, the enhanced MBP expression we observed in all hippocampal subregions of Kir4.1 ko mice was unlikely to be due to onset of myelination of former NG2 glia. Rather, lack of Kir4.1 in NG2 glia might have stimulated myelination in already existing oligodendrocytes. Indeed, adaptive myelination and MBP expression depend on neuronal activity and neuron-glia interactions (Bechler et al., 2018; ***Timmler and Simons***, 2019) and NG2 glia contacting nodes of Ranvier may sense the status of myelination (***Serwanski et al***., 2017). MBP expression and myelination in male brains exeeds that in females (***Cerghet et al***., 2006; ***Markham et al***., 2009), and sex differences also exist in hippocampal learning and morphology (***Koss and Frick***, 2017). Notably, however, we have observed differences between male and female myelination and behavior only after deletion of Kir4.1, but not in wildtype mice.

Grey matter myelination continues throughout adulthood and includes both, formation of new and plasticity of already established myelin internodes (***Hill et al***., 2018; ***Hughes et al***., 2018). To further evaluate how deletion of Kir4.1 from NG2 glia affected myelination, expansion microscopy was performed allowing for higher (factor 4-5) spatial resolution than conventional confocal microscopy (***Chen et al***., 2015; ***Chozinski et al***., 2016). MBP is only one among several myelin components, however its strong impact on myelin compactness (***DeBruin et al***., 2005) makes it an useful measure to assess myelination. We found increased density of MBP in the myelin sheaths as a consequence of targeted deletion of Kir 4.1. Possibly, enhanced Ca^2+^ signaling in oligodendrocytes lacking Kir4.1 underlies enhanced MBP expression and myelin formation (***Zhang et al***., 2019).

### Impaired LTP in the hippocampus of mice with Kir4.1 deficient NG2 glia

Deletion of Kir4.1 from NG2 glia not only affected neuron-NG2 glia communication but also altered neuronal network function and animal behavior. Basal excitability of CA1 neurons and vesicle release probability were not affected, but field potential recordings at the CA3 - CA1 synapse revealed impaired LTP in mice lacking Kir4.1 in NG2 glia indicating functional alterations at the postsynaptic site. Conditional deletion from all macroglia resulted in enhanced LTP at the same synapse (***Djukic et al***., 2007), presumably due to impaired K^+^ and glutamate homeostasis, i.e. functions through which astrocytes regulate neuronal excitability (***Katagiri et al***., 2001; ***Wallraff et al***., 2006; ***Filosa et al***., 2009; ***Chever et al***., 2010; ***Sibille et al***., 2014). In the somatosensory cortex, shedding of the NG2 ectodomain is required for proper LTP, probably by facilitating expression of Ca^2+^ impermeable AMPA receptors (***Sakry et al***., 2014). However, deletion of NG2 did not affect LTP in the hippocampus (***Passlick et al***., 2016). The secretome of NG2 glia is not yet well characterized. Neurotrophic factors bFGF and FGF-2 are released by astrocytes, neurons and microglia cells and influence neuronal plasticity, learning and memory and also LTP at CA3 - CA1 synapses (***Terlau and Seifert***, 1990; ***Zechel et al***., 2010). In a mouse model of chronic stress disorder, loss of NG2 glia was found in the CA1 region, and it was suggested that reduced release of FGF-2 from NG2 glia causes loss of astrocyte and neuronal function and depressive behavior (***Birey et al***., 2015). Thus, it is conceivable that in our mice with targeted deletion of Kir4.1, impaired release of growth factor(s) from NG2 glia affected hippocampal LTP.

### Behavioural changes

Reduced LTP after high frequency stimulation of Schaffer colaterals in hippocampal slices of Kir4.1 ko mice did not entail deficits in behavioral tests. Instead, mice with targeted deletion of Kir4.1 in NG2 glia showed improved declarative memory in the NOLR test. A negative correlation of CA3-CA1 LTP and behavior, i.e. enhanced LTP but impaired behavioral performance, was previously observed after genetic deletion of transcription activator Fmr2 (***Gu et al***., 2002), PSD-95 (***Migaud et al***., 1998) or heterozygous deletion of the BDNF receptor trkB (***Minichiello et al***., 1999). However, homozygous trkB receptor knockout caused decreased LTP and behavioural impairment suggesting that only profound LTP reduction affects behavior (***Qi et al***., 1996; ***Minichiello et al***., 1999). Thus, the improved performance in the NOLR test of our NG2 Kir 4.1 ko mice probably resulted from increased myelin compactness. Indeed, several previous studies have reported that myelination deficits are accompanied by impaired cognitive performance (***Geraghty et al***., 2019), social isolation (***Makinodan et al***., 2012) or deficits in spatial memory and motor skills (***McKenzie et al***., 2014; ***Xiao et al***., 2016; ***Steadman et al***., 2020). Myelin ensheathment and MBP compactness coordinate temporal regulation and synchronicity of neuronal firing and circuit function (***Salami et al***., 2003; Seidl, 2014) as well as speed of action potential propagation (***Min et al***., 2009), and its impairment results in dispersed synaptic transmission, asynchronous firing and deficits in motor learning.

Notably, improved performance of Kir4.1 ko mice was confined to the NOLR test, while no differences were found in working memory (Y-maze) or social memory (partner recognition test). It has to be considered that not only the hippocampus is involved in executing these memory types. The prefrontal cortex contributes to short term memory performance in the Y-maze test (***Lynch***, 2004; ***Albayram et al***., 2016) and other areas of the limbic system, e.g. the amygdala, play a role in partner recognition (***Kogan et al***., 2000; ***Bilkei-Gorzo et al***., 2014). For NOLR, which is based on information about the object itself (familiar or new) and its location, the medial temporal lobe including perirhinal, entorhinal and parahippocampal cortices, are important (***Squire and Dede***, 2015). Thus, the changes occurring in the hippocampus upon deletion of Kir4.1 in NG2 glia affect memory performance differently, with some behavioral tasks being more susceptible to the deletion-induced alterations than others, but hippocampal memory deficits and motor deficits were not observed.

## Conclusions

Activity-dependent myelination is an important mechanism to ensure proper synaptic function and spiking synchronicity, motor skills and memory performance. Our data imply that these processes are also influenced by NG2 glia and that this is not simply dependent on their differentiation status. Rather, it is likely that NG2 glia exert its effect by signaling on neighboring oligodendrocytes, either directly or indirectly via influencing neuronal activity. Future work has to identify the mechanisms and factors by which NG2 glia accomplish these actions.

## Materials and Methods

Experiments were performed with male and female NG2ki-EYFP (***Karram et al***., 2008) and NG2-CreERT2 x Kir4.1 fl/fl x Rosa26-EYFP mice. The latter were obtained by cross breeding mice carrying the floxed Kir4.1 gene (Kir4.1 fl/fl) with knockin mice expressing the Cre DNA recombinase variant CreERT2 in the NG2 (*cspg4*) locus (NG2-CreERT2 x Rosa26-EYFP) as previously described (***Huang et al***., 2014). NG2-CreERT2 x Rosa26-EYFP mice (without floxed Kir4.1) were used as controls. In both transgenic lines, Rosa26-floxed-stop-EYFP (Rosa26-EYFP) is used as a reporter to monitor Cre activity (***Srinivas et al***., 2001; ***Huang et al***., 2014). Mice were kept under standard housing conditions conditions (12/12h light/dark cycle, food and water *ad libitum*). All experiments were carried out in accordance with local, state and European regulations (veterinary licenses 84-02.04.2015.A411, 81-02.04.2017.A437).

### Tamoxifen administration

Mice of either sex (p23-26) were intraperitoneally injected with 1.5 mg tamoxifen (Sigma) dissolved in ethanol (AppliChem, Roth) and sunflower seed oil (Sigma) in a ratio of 1:10. Animals were injected once per day for three consecutive days (***Huang et al***., 2014). Electrophysiological and immunohistochemical analyses were performed 3-4 and 8 wpi, respectively. For the behavioral experiments the mice were introduced to the tasks in the 3rd week after tamoxifen.

### Slice preparation

Mice were anesthetized with 50% O_2_/50% CO_2_ or isoflurane before decapitation. Horizontal slices of 200-300 µm thickness were obtained with a vibratome (Leica VT1000S/1200S; Leica Microsystems) in ice cold slicing solution containing (in mM): 87 NaCl, 2.5 KCl, 1.25 NaH_2_PO_4_, 7 MgCl_2_, 0.5 CaCl_2_, 25 NaHCO_3_, 25 glucose, 61.3 sucrose. Sections were incubated for 15 min at 35°C in the same solution before being stored at room temperature in artificial cerebrospinal fluid (aCSF; in mM: 132 NaCl, 3 KCl, 2 MgCl_2_, 2 CaCl_2_, 10 glucose, 1.25 NaH_2_PO_4_, 20 NaHCO_3_) until further use. For extracellular field recordings, the bath solution contained (in mM) 131 NaCl, 2.5 KCl, 1.3 MgSO_4_, 1.25 NaH_2_PO_4_, 21 NaHCO_3_, 2 CaCl_2_, 10 glucose. Solutions were oxygenated with carbogen (5% CO_2_/95% O_2_). Slices were allowed to recover for at least 1 h before experiments.

### Patch-clamp recordings

Slices were transferred to a recording chamber on an upright microscope (Axioskop FS2, Zeiss, equipped with a CCD camera, VX45, Optronics, Infrared-DIC optics, Eclipse E600 FN, Nikon, and epifluorescence, Polychrome II, Till Photonics), and constantly perfused with oxygenated aCSF. NG2 glial cells located in the hippocampal CA1 stratum radiatum were identified by intrinsic EYFP expression after induction of Cre mediated recombination by tamoxifen. Borosilicate capillaries (Science Products) pulled by a horizontal puller (P-87; Sutter Instrument) had a resistance of 3-5 MΩ when filled with either of the following intracellular solutions (in mM): 125 K-gluconate, 20 KCl, 3 NaCl, 2 MgCl_2_, 0.5 EGTA, 3 Na_2_-ATP, 10 HEPES (for miniature EPSP (mEPSP) recordings; liquid junction potential corrected) or 130 KCl, 2 MgCl_2_, 0.5 CaCl_2_, 3 Na_2_-ATP, 5 BAPTA, 50 µM spermine, 10 HEPES (for evoked EPSC recordings). Cells were clamped to -80 mV, and series and input resistance were constantly monitored applying 10 mV voltage steps. Whole cell patch-clamp recordings were obtained using an EPC800 amplifier (HEKA Electronic) and digitized with an ITC 16 D/A converter (HEKA Electronic). Signals were filtered at 1 or 10 kHz, sampled at 6, 10 or 30 kHz and displayed by Tida software (HEKA Electronic). In the current-clamp mode, voltage signals were amplified with a DPA 2FS amplifier (npi electronic). Experiments were performed at 35°C if not stated otherwise.

Passive membrane properties were determined after establishing the whole-cell configuration. The average of 10 consecutive transients evoked by 50 ms lasting 10 mV voltage step was used to determine series resistance (Rs), membrane resistance (Rm) and membrane capacitance (Cm). As it is currently impossible to properly determine resistances above 3 GΩ due to technical reasons, data above this threshold was excluded from analysis. mPSPs were recorded (8 min periods) in aCSF containing TTX (0.5 µM; Abcam). NBQX (10 μM; Tocris) and/or Picrotoxin (150 µM; Abcam) were applied via the perfusion system to separate GABAergic and glutamatergic inputs. mEPSPs were detected using the template search of pClamp (Molecular devices) and evaluated by visual inspection. Cells with less than 3 events detected were rejected. Amplitudes, kinetics and frequencies of mEPSPS were analyzed by a custom-written macro in IGOR Pro Software (WaveMetrics).

### Schaffer collateral stimulation

Schaffer collaterals were stimulated using a monopolar stimulation electrode consisting of a teflon-coated silver wire and a low-resistance glass pipette (<1 MΩ) filled with aCSF. The pipette was placed in the CA1 stratum radiatum and stimulus intensity was adjusted to stimulate single or few presynaptic fibers (minimal stimulation). Paired pulse stimulation was induced by biphasic voltage pulses of 150 µs duration and inter-stimulus intervals of 50 ms, applied via an AM-Systems isolation pulse stimulator (Model 2100, A-M Systems). Recombined NG2 glial cells were recorded with KCl-based intracellular solution. A response was considered a failure if the current amplitude was less than twice the baseline noise, verified by visual inspection. Recordings were analyzed by a custom-written macro in IGOR Pro Software (WaveMetrics).

### Field potential recordings

Experiments were performed in an interface chamber (IFC) (custom-made, AG Heinemann, Charité-Universitätsmedizin Berlin), and slices were continuously perfused with carbogenated aCSF (34.0 ± 0.5°C; flow rate, 2.5 ± 0.2 ml/min). For stimulation, a concentric bipolar electrode was used (CBARC75; FHC) and stimulus intensity was controlled by an isolated current stimulator (DS3, Digitimer Ltd.). Signals were pre-amplified, filtered (highpass 0.1 Hz, lowpass 20 kHz; EXT-02B, npi) and 50 Hz-interferences were removed (HumBug, Quest Scientific Instruments Inc.). Signals were sampled at a frequency of 10 kHz (NI USB-6221, National Instruments). Hippocampal evoked field potentials were recorded in the CA1 stratum radiatum using patch pipettes (resistance 3-8 MΩ) filled with aCSF. Input-output curves were obtained with increasing stimulation intensities (range: 20 - 500 μA). Each stimulus intensity was delivered 3 times, mean values were calculated and the initial slope of the fEPSPs was evaluated. Baseline paired-pulse (inter-stimulus interval 50 ms) or single pulse (100 µs half-wave pulse duration) recordings were obtained every 15 s for 10 min. LTP was induced by theta burst stimulations (TBS), applying 8 stimulus trains at 5 Hz (4 pulses at 100 Hz; 1 min inter-stimulus interval, 3 repetitions). After TBS, baseline stimulation was repeated for further 30 min (120 pulses). Stimulation intensity was adjusted to evoke half-maximal fEPSP amplitudes. Analysis was performed with Clampfit (V. 10.3; Molecular Devices, LLC). fEPSPs elicited by paired-pulse stimulation were normalized to the mean fEPSP slopes obtained during the first 10 min of the experiment (baseline). To evaluate LTP, responses recorded before (baseline, 10 min) and after TBS were normalized to the mean fEPSP slope during baseline recording and the ratio of fEPSP slopes was calculated (paired-pulse ratio= slope 2 / slope 1). Recordings with more than 10% fEPSP slope variability during baseline recording were excluded.

### FACsorting of NG2 glial cells

Mice of both sexes (NG2ki-EYFP at p60; Kir4.1 ko and NG2CreERT2xRosa26-EYFP (control) mice 3wpi) were sacrificed, their brains were dissected, and whole hippocampi were isolated under microscopic control (Stereo microscope, Zeiss, Germany). Cell suspensions were prepared by mincing and digesting the tissue in papain (37 °C, 15 min). Then, DNAseI was added (incubation for another 10 min; Neural Dissociation Kit (P), Miltenyi, Germany). The tissue was dissociated by Pasteur pipettes, filtered through a 70 µm cell strainer and centrifuged (300 x g, 10 min) after addition of 10 ml HBSS (with Ca^2+^ and Mg^2+^). The pellet was re-suspended in 1 ml HBSS (without Ca^2+^ and Mg^2+^), filtered through a 40 µm cell strainer. NG2 cells were identified by their fluorescence (emission at 527 nm) and sorted by a FACSAriaIII flow cytometer (70 µm nozzle, BD Biosciences, Heidelberg, Germany) into tubes in HBSS (without Ca^2+^ and Mg^2+^). After centrifugation at 2,000 x g (10 min) the supernatant was discarded, the cells were re-suspended in 200 µl lysis/binding buffer (Invitrogen, Darmstadt, Germany), frozen in liquid nitrogen and stored at -80 °C.

### Semiquantitative real-time RT-PCR

Messenger RNA was isolated from isolated cells by cell lysis in the lysis/binding buffer and by using oligo(dT)25-linked Dynabeads (Invitrogen). The beads with adherent mRNA were suspended in DEPC-treated water (20 µl). For first strand synthesis, first strand buffer (Invitrogen), dithiothreitol (DTT, 10 mM), dNTPs (4x 250 µM, Applied Biosystems), oligo-dT_24_-primer (5 µM, Eurogentec), RNasin (80 U, Promega) and SuperscriptIII reverse transcriptase (400 U, Invitrogen) were added and the reaction mix was incubated for 1 h at 50 °C (final volume 40 µl). The reaction mixture for real-time PCR contained Takyon real-time PCR mastermix (Eurogentec) and Taqman primer/probe mix (Applied Biosystems, Darmstadt, Germany), and 1 µl of the RT-product was added (reaction volume 12.5 µl). PCRs for the respective target genes and β-actin as a housekeeping gene were run in parallel wells for each sample, respectively and triplicates for each sample were performed. Negative controls (water) were also performed in each run. Samples were incubated at 50 °C (2 min), and after denaturation (95 °C, 10 min), 50 cycles were performed (denaturation at 95 °C, 15 s; primer annealing and extension at 60 °C, 60 s). Fluorescence intensity was readout during each annealing/extension step (CFX 384, Biorad, Munich, Germany). The target gene/β-actin gene expression ratio was determined by comparing C_T_ values of the target gene with those of the reference gene. The relative quantification of different genes was determined according to the following equation:

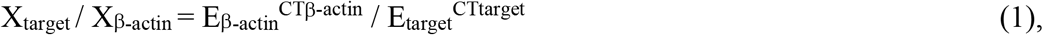

yielding the gene ratio with X being the input copy number, E the efficiency of amplification, and C_T_ the cycle number at threshold. By quantification of target gene expression against that of β-actin, C_T_ was determined for each gene at the same fluorescence emission, R_n_. The amplification efficiency was determined by serial dilutions of mRNA and was 1.93 for Kir4.1, 1.94 for β-actin, 1.91 for MAG and 1.89 for MBP.

### Single-cell RT-PCR

Single-cell transcript analysis was performed as previously reported (***Matthias et al***., 2003). Briefly, after recording, the cytoplasm of individual cells was harvested under microscopic control (Axioskop FSII, Zeiss, equipped with a CCD camera VX45, Optronis, Kehl, Germany) and aspirated in reaction tubes filled with 3 µl DEPC-treated water. Reverse transcription (RT) was performed after addition of first strand buffer (Invitrogen), dithiothreitol (DTT; 10 mM), dNTPs (4x 250 µM), random hexamer primers (50 µM; Roche), RNasin (20 U; Promega) and SuperscriptIII reverse transcriptase (100U; Invitrogen) at 37 °C for 1h. A multiplex two-round single-cell PCR was performed with primers for Kir4.1 and PDGFRα (Table 1). The first PCR was performed after adding PCR buffer, MgCl_2_ (2.5 mM), primers (200 nM each), and 5 U *Taq* polymerase (Invitrogen, Karlsruhe, Germany) to the RT product (final volume 50 µl). Forty-five cycles were performed (denaturation at 94 °C, 25 s; annealing at 51 °C, 2 min for the first 5 cycles, and 45 s for the remaining cycles; extension at 72 °C, 25 s; final elongation at 72 °C, 7 min). An aliquot (2 µl) of the PCR product was used as template for the second PCR (35 cycles; annealing at 54 °C, first 5 cycles: 2 min, remaining cycles: 45 s) using nested primers (Table 1). The conditions were the same as described for the first round, but dNTPs (4x50 µM) and Platinum *Taq* polymerase (2.5 U; Invitrogen) were added. Products were identified with gel electrophoresis using a molecular weight marker (Low molecular weight marker, New England Biolabs, Frankfurt, Germany). As a positive control, RT-PCR for total RNA from mouse brain were run in parallel. Negative controls were performed using distilled water or bath solution for RT-PCR. The primers for the targets were located on different exons to prevent amplification of genomic DNA.

**Table 1.** Membrane properties of control, Kir4.1 wt and Kir4.1 ko cells. Data is given as median and interquartile range (quartile 25% – quartile 75%), number of cells in square brackets. RMP, resting membrane potential; Rm, membrane resistance; Cm, membrane capacitance.

|  | **Control** | **Kir4.1 wt** | **Kir4.1 ko** |
| --- | --- | --- | --- |
| **RMP (mV)** | -89<br>(-90 - -86)<br>[82] | -85<br>(-89 - -82)<br>[55] | -61<br>(-69 - -53)<br>[128] |
| **Rm (MΩ)** | 50.75<br>(32.69 – 81.75)<br>[82] | 81.22<br>(56.56 – 115.17)<br>[55] | 546.10<br>(348.94 – 1022.90)<br>[118] |
| **Cm (pF)** | 29.98<br>(25.68 – 35.66)<br>[82] | 31.09<br>(24.92 - 38.16)<br>[55] | 27.28<br>(23.09 – 33.89)<br>[128] |
| **Conductance at -130 mV (pA/mV)** | 17.42<br>(13.78 – 22.62)<br>[82] | 12.81<br>(9.77 – 17.46)<br>[55] | 1.40<br>(0.93 – 2.05)<br>[128] |

### Immunohistochemistry

Mice were anesthetized by intraperitoneal injection of Xylaxin/Ketamin and intracardially perfused by ice cold phosphate buffer saline (PBS) followed by 4% paraformaldehyde (PFA). Brains were dissected, post fixed overnight in 4% PFA at 4°C and transferred into PBS. Coronal brain sections (40 µm thickness) were obtained with a vibratome (Leica VT1200S; Leica Microsystems) and washed three times with PBS before incubating in blocking solution (0.5% Triton-X100 and 10% normal got serum (NGS) in PBS) for 2h at room temperature. We used the following primary antibodies (diluted in PBS with 0.1%Triton-X100 and 5%NGS, applied for at least 12 h at 4°C): chicken anti-GFP (1:500; Abcam), rabbit anti-MBP (1:250; Millipore), mouse anti-GSTpi (1:500; BD Bioscience), rabbit anti-Ki67 (1:500; Novocastra), rabbit anti-PDGFRα (1:200; Thermo Fisher). Secondary antibodies: Goat anti-chicken Alexa 488 (1:500; Thermo Fisher), goat anti-rabbit Alexa 594 (1:500; Thermo Fisher), goat anti-rabbit Alexa 647 (1:500; Thermo Fisher), goat anti-mouse Alexa 647 (1:500; Thermo Fisher) (all in PBS with 0.1%Triton-X100 and 5% NGS, 2h at room temperature). After nuclei staining with Hoechst (Molecular Probes, 10 min, 1:100 in dH_2_O) sections were washed again and mounted on cover slips (Aqua-Poly/Mount, Polyscience).

### Confocal microscopy

Images were captured with a confocal laser scanning microscope (SP8 LSM, Leica) using a 20x immersion objective (Leica). Z-stack images of 1 µm interval (total depth of 4 µm) were processed with LAS software (Leica) and analyzed with Fiji software. Images were acquired and analyzed from 3 hippocampal sections per mouse, from at least 6 animals per group. To evaluate recombination, differentiation, proliferation and myelination, the data from strata radiatum, lacunosum moleculare and moleculare of the DG were averaged, accordingly. Recombination efficiency was calculated as the proportion of cells co-expressing EYFP and PDGFRα among all PDGFRα^+^ cells. To assess changes in the differentiation of recombined (i.e. EYFP^+^) NG2 glia cells into oligodendrocytes, sections were stained for the oligodendrocyte marker GSTpi and the proportion of cells co-expressing EYFP and GSTpi among all EYFP^+^ cells was determined. Proliferation was quantified by calculating the ratio of cells co-expressing Ki67 and EYFP among all EYFP cells. For quantification of myelination, the mean grey intensity (calculated as the integrated fluorescence intensity per area) was measured and background subtracted. All data were normalized to the appropriate staining of control sections from NG2-CreERT2 x Rosa26-EYFP mice.

### Expansion Microscopy

The expansion microscopy (ExM) protocol for imaging proteins with conventional primary and secondary antibodies was adopted from ***Deshpande et al***. (2017); ***Herde et al***. (2020) and ***Asano et al***., (2018). Fixed coronal hippocampal slices (70 µm thickness) were cut on a vibratome and blocked overnight at 4°C in blocking buffer (5% normal goat serum, 1% Triton-X100 in PBS pH 7.4). Primary antibodies were incubated in blocking buffer for 48h at 4°C. Antibodies used were: chicken anti-GFP (1:1000; Abcam ab13970, lot: GR236651-g), rabbit anti-MBP (1:200; Millipore AB980, lot: 2869285) and mouse anti-SMI312 (1:100; Biolegend 837904, lot: B263754). After washing (PBS, 3x20 min, RT), secondary antibodies were incubated overnight at 4°C in blocking buffer. Secondary antibodies used were: goat anti-chicken Alexa Fluor 488 (1:200; ThermoFisher A11039, lot: 1691381), goat anti-mouse Alexa Fluor 568 (1:200; ThermoFisher A1131) and goat anti-rabbit biotin (1:200; Jackson ImmunoResearch 111-066-144). The following day, after washing (PBS, 3x20 min, RT), Hoechst 33342 was incubated for 10 min at RT in distilled water (1:2000). After washing (PBS, 3x20 min, RT), the slices were pre-imaged with 40x/1.1 NA objective in a Leica SP8 confocal microscope and a z-stack (1 µm interval) of the tip of the DG acquired for expansion factor calculation (only Hoechst 33342 signal imaged). Further treatment was adopted from ***Deshpande et al.***, (2017) and ***Asano et al***., (2018). Briefly, slices were incubated in 1 mM methylacrylic acid-NHS for 1h at RT. After washing (PBS, 3x20 min, RT), slices were incubated for 45 min at 4°C in monomer solution (in g/100 mL PBS: 8.6 sodium acrylate, 2.5 acrylamide, 0.15 N,N’-methylenebisacrylamide, 11.7 NaCl). Then, slices were incubated for 5 min at 4°C in gelling solution (monomer solution supplemented with %(w/v): 0.01 4-hydroxy-TEMPO; 0.2 TEMED; 0.2 ammonium persulfate). Slices in the gelling solution were placed on a glass slide and the preparation was covered with a coverslip and incubated for 2h at 37°C. To ensure that all samples have the same thickness, the gel was sandwiched between a glass slide and a coverslip, with precise spacers at the edges with 170 ± 5 µm thickness. Coverslips and excess gel around the slice were removed after gelification and gels incubated overnight at 25°C in digestion buffer containing (50 mM Tris pH 8.0, 1 mM EDTA, 0.5% Triton-X100, 0.8 M guanidine hydrochloride, 16 U/ml of proteinase K). The following day, the digestion buffer was removed and the gels were incubated with streptavidin Alexa Fluor 647 (1:100; Jackson ImmunoResearch 016–600-084, lot: 124695) in PBS for 2h at RT. For expansion, slices were then incubated in distilled water (adjusted pH 7.4 with NaOH) for 2.5h at RT and water exchanged repeatedly every 15-20 min. Finally, slices were mounted on poly-lysine coated µ-Slide 2 well Ibidi-chambers and sealed with a poly-lysine coated coverslip on top, adding a drop of water to prevent the gel from drying (ibidi chambers and coverslips were poly-lysine coated by incubation with poly-l-lysine solution (0.1% w/v in water (P8920, Sigma-Aldrich, lot: 050M4339) for at least 45 min at RT shaking and dried with pressured air). Prior to imaging the region of interest (ROI) for each slice, the tip of the DG was imaged with an 20x/0.75 NA objective (z-stack 6 µm interval with Hoechst 33342 signal only). The expansion factor was determined by identifying the same cells labeled with Hoechst 33342 in the tip of DG before and after expansion and averaging the expansion factor of 10 measures. On average, we obtained an expansion factor of 4.546 ± 0.242 (n = 10). ROIs were imaged on a Leica SP8 inverted confocal microscope using a 40x/1.1 NA objective and hybrid detectors. Image stacks were then deconvolved in Leica Systems software and analyzed with Fiji. 3D-Isosurfaces were constructed for MBP and SMI312 immunofluorescence using IMARIS 8.31 (Bitplane, Zurich, Switzerland) isosurface routines including intensity thresholding, background correction and minimal object volume. This set of parameters was optimizes for best fitting result, verified by visual inspection and kept constant for all analyses. Volume, summed fluorescence intensity and fluorescence density of the individual 3D objects generated in this way were determined for 4 different ROIs for control and Kir4.1 ko animals. Physical xyz-voxel size (i.e. after expansion) of confocal stacks of interest was 0.284 x 0.284 x 0.8 µm^3^ taken as 1024 x 1024 x 60 pixels. This corresponds to an original (i.e. before expansion) ROI size of 66.8 x 66.8 x 11 µm^3^

### Behavioral tests

Behavioral tests were performed as previously reported (***Albayram et al***., 2016; ***Bilkei-Gorzo et al***., 2017). The beam walking test was carried out in a sound isolated and well-lit chamber, where a 1 m long rod was fixed to a shelf at the height of 70 cm. An escape chamber (made from dark plastic, 7 x 10 x 10 cm^3^, its floor covered with saw dust) was positioned at the end of the rod. On day 1, mice were placed to the free end of a rod with 28 mm diameter and were allowed to climb to the escape box. Mice, which fell down were placed back to the original position. After entering the escape (goal) chamber mice were allowed to rest for a minute and then the test was repeated 2 additional times. At the end of this period each animal readily and quickly walked from the start position to the goal chamber. On day 2, the test was carried out again as described above, but in this case we measured the time from starting until entering (with all 4 legs) into the goal box (latency). All animals were tested first using the thickest (28 mm diameter), next the middle sized (14 mm) and finally the thinnest (8 mm) rod.

For the Y-maze test, the length of the arms of the Y-shaped labyrinth was 20 cm and the height of their walls was 15 cm. Animals were placed into the end of one of the arms and their location and activity was recorded and analysed by the EthoVision tracking system (Noldus) for 10 min. The ration of spontaneous alternations was defined as entering a different arm in 3 consecutive arm entries. Percent value was calculated as SA/TA-2 *100, where SA is the number of spontaneous alternations and TA is the total number of arm entries. The maze was thoroughly cleaned using detergent between the animals to avoid odour cues.

The novel object location recognition test was performed in a sound-isolated, dimly illuminated room in an open field-box (44 x 44 cm^2^). The floor was covered with sawdust (1 cm deep, saturated with the odour of the animals during habituation). The habituation period consisted of a daily 5-min period of free exploration in the arena for 3 days. On day 4, the animals were allowed to explore 3 identical objects (LEGO pieces with different colours, roughly 2 x 2 cm^2^) placed into the area in a fixed location for 6 min, and the time spent on the inspection of the individual objects was recorded (Noldus Ethovision, XT software). Thirty min later the animals were placed back into the box, where one object was placed into a new location. The animals were left to explore for an additional 3 min. The time spent with investigations was recorded, and the preference ratio for the moved object was calculated as follows: Preference= Ta/(Ta+Tb+Tc)*100, where T is the time spent with investigation, a – object which is moved in the second trial, b and c objects remained in the original position. Novelty preference was calculated as (Pt2-Pt1)/Pt1*100, where P is the preference, t1 is trial 1 and t2 is trial 2.

The partner recognition test was performed in the same arenas and after the same habituation as described for the novel object location recognition test. In the first trial, the arenas contained a metal grid cage, with a mouse in one cage (same age and sex as the test animal, from different cage) and an object (similar size and form as the metal grid cage) in the opposing corner 6-7 cm from the walls. Location and activity of the test mouse were recorded (15 min) and analyzed (EthoVision tracking system, Noldus). In the next trial, the object was replaced with another grid cage containing a new partner and the activity of the test mouse was recorded again (5 min). The inter-trial interval was 1 h on day 4, whereas 2, 4 and 8 h on day 5, 6 and 7, respectively. Recognition of the previously seen partner was defined by a novelty preference i.e. a significantly higher period spent investigating the new partner in the second trial. Novelty preference was calculated as (Ta-Tb)/Tb*100, where Ta is the time spent with the novel, and Tb is the time spent with the previous partner.

### Statistics

Data was analyzed with Igor Pro (WaveMetrics), (R Core Team (2020) and Origin (OriginLab) software. All data was tested for normal (Gaussian) distribution (Shapiro–Wilk test or Lilliefors test, dependent on the number of data points), followed by two sampled Student’s t or Mann-Whitney U test with or without Welch correction for equal or diverse variances. For group analysis data was tested with Kruskal–Wallis ANOVA followed by Dunńs test, 1-way ANOVA followed by Tukey test or 2-way ANOVA followed by Bonferronis t-test. Significance level was set to P<0.05. Non-Gaussian distributed data is displayed in Tukey box plots showing median (central line), quartiles (25% and 75%; box) and whiskers (± 1.5 times the interquartile range). Outliers are shown as dots. A density estimator with a Gaussian kernel approximated empiric distributions to comparable probability density functions. We used the sm package of R for this purpose.

## Acknowledgements

We thank T. Erdmann for technical assistance.

---

**Suppl. Figure 1:**
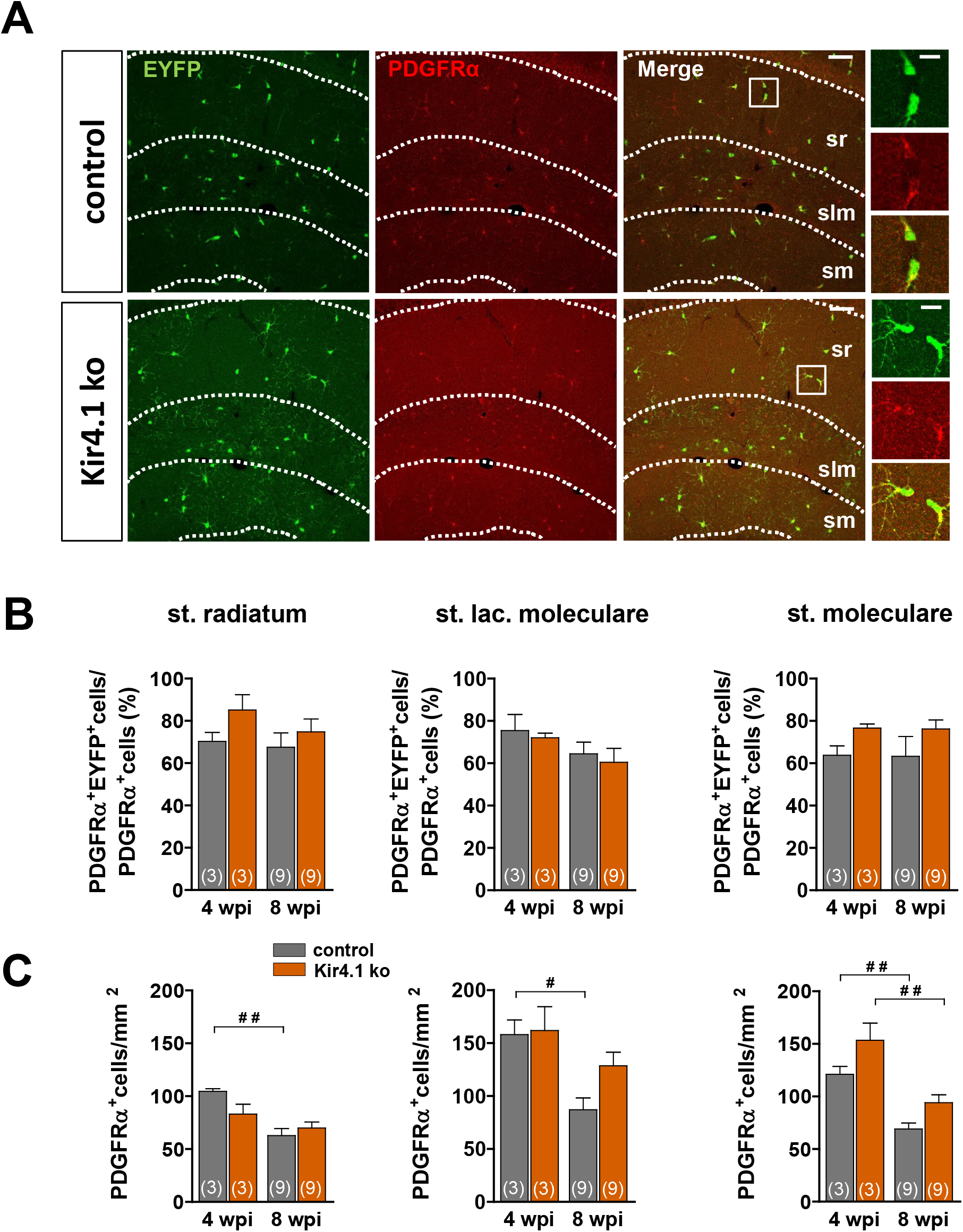
Assessing recombination efficiency of Kir4.1 deletion in NG2 glia by antibody staining, 4 and 8 weeks after tamoxifen injection. A) Immunostainings for EYFP (green) and the NG2 glia marker, PDGFRα (red), in hippocampal slices from control and Kir4.1 ko mice. Blow-ups of boxed areas are given to the right. Scale bar 60 and 15 µm (blow-ups). B) Recombination efficiency was determined by dividing the number of EYFP^+^PDGRα^+^ cells by the number of PDGRα^+^ cells. Four and 8 wpi of tamoxifen, 60-80% of all NG2 glia expressed EYFP, indicating recombination. C) The density of PDGFRα^+^ cells was similar between control and Kir4.1 ko mice, 4 and 8 weeks after injection. However, in control mice, the density of PDGFRα^+^ cells decreased over time in all three layers (st. radiatum, p = 0.008; st. lac. moleculare, p = 0.033; st. moleculare, p = 0.005) while in Kir4.1 ko mice, a decrease was only seen in the stratum moleculare of the DG (p = 0.002). 2-way ANOVA post-hoc Tukey test, mean ± SEM. Number of mice is given in bar graphs. # p<0.05, ## p< 0.01.

**Suppl. Figure 2:**
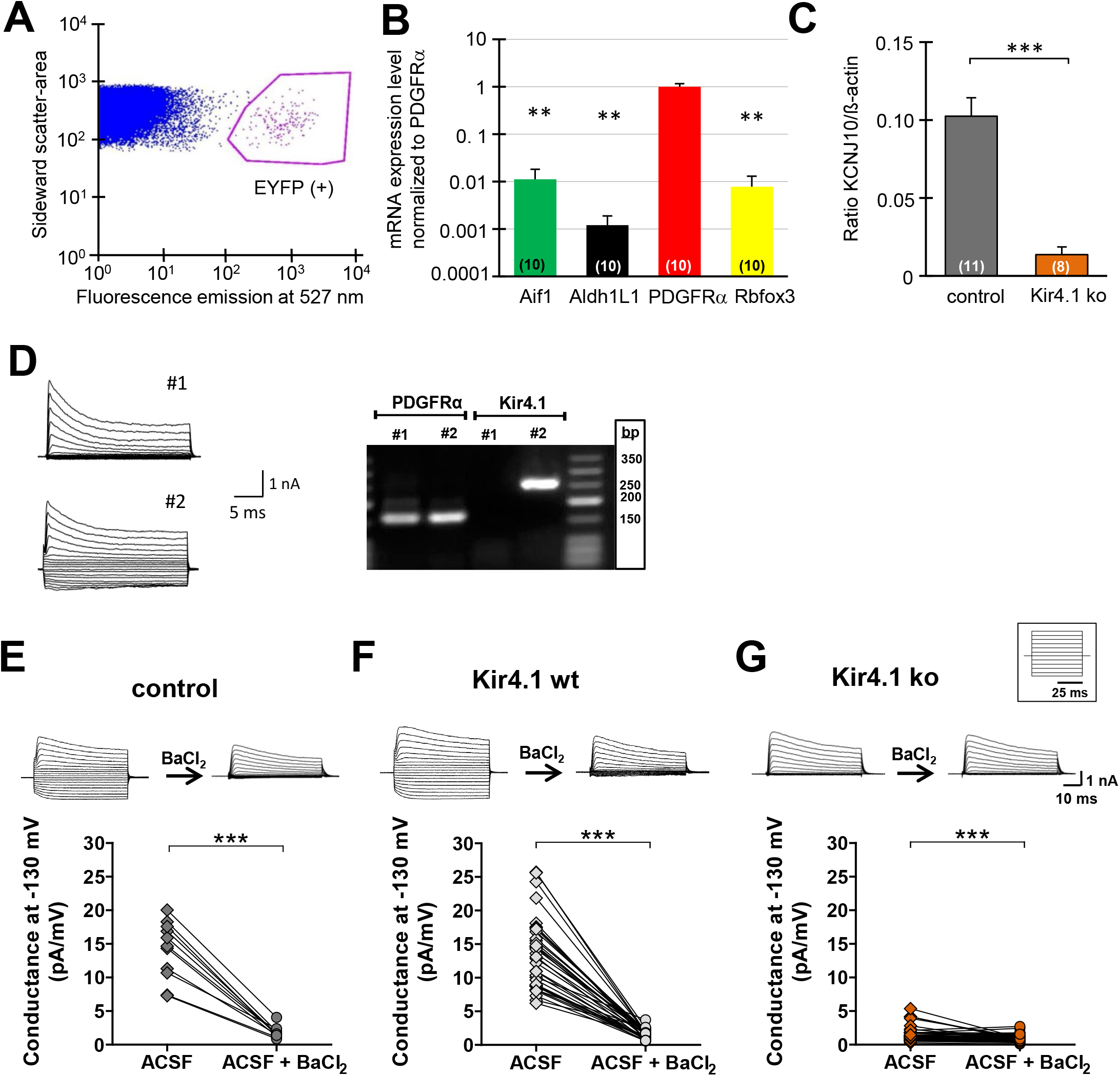
Assessing recombination efficiency of Kir4.1 deletion in NG2 glia by transcript analyses and the Ba^2+^ sensitivity of inward currents. (A) Scatter plot of FAC sorted recombined NG2 glia (EYFP+) of the hippocampus from control and Kir4.1 ko mice. (B) Semi quantitative PCR confirmed specificity of FAC sorted NG2 glia (PDGFRα) from the hippocampus. Gene transcripts of astrocytes (Aldh1L1), neurons (Rbfox3) and microglial cells (Aif1) were almost absent (n = 10). ** p < 0.05, T-test, mean ± SEM. Number of mice is given in bar graphs. (C) Reduction of Kir4.1 mRNA expression to about 15% in recombined NG2 glia in the hippocampus. Number of mice is given in parentheses. *** p < 0.001, T-test, mean ± SEM. Number of mice is given in bar graphs D) Single-cell RT-PCR was performed with recombined NG2 glia harvested from slices of Kir4.1 ko mice. Cells were tested for the presence of mRNA for PDGFRα and Kir4.1. In NG2 glia lacking inward currents (Kir4.1 ko, cell #1; conductance at -130 mV: <6pA/mV), absence of Kir4.1 mRNA was confirmed. In recombined NG2 glial cells where inward currents could still be elicited, Kir4.1 mRNA was still present (Kir4.1 wt, cell #2). E-G) Whole-cell currents of NG2 glia were evoked applying de- and hyperpolarizing steps between -160 mV and +20 mV, before and after focal pressure application of the Kir channel blocker BaCl_2_ (100 μM; holding potential -70 mV). The membrane conductance was calculated at -130 mV, which decreased in the presence of BaCl_2_ (control cells, n = 11; Kir4.1 wt, n = 35; Kir4.1 ko, n = 49). Membrane conductance of individual cells is indicated before (diamonds) and during (circles) BaCl_2_ application.

**Suppl. Figure 3:**
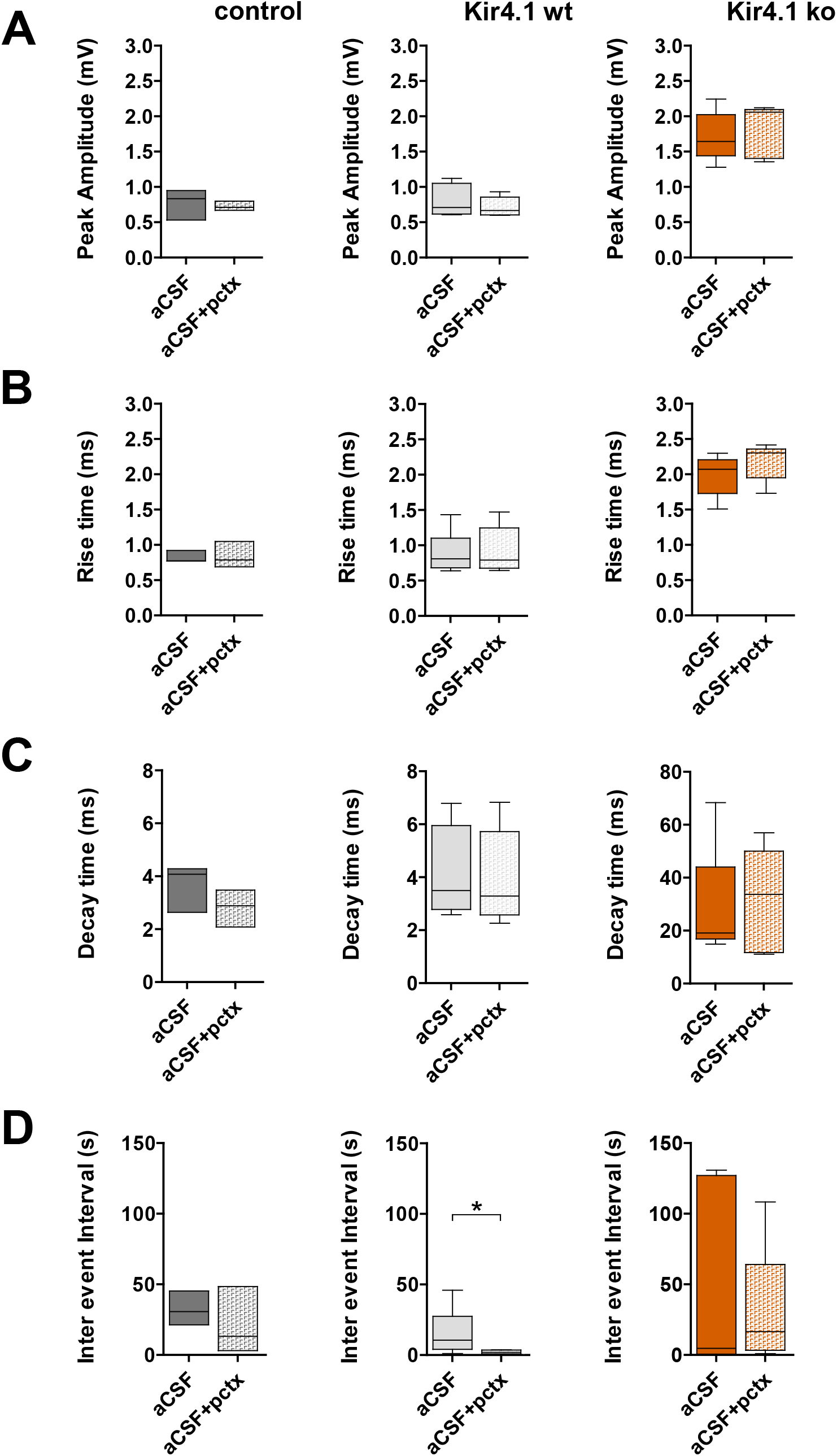
Picrotoxin does not affect mPSPs of NG2 glia. A-C) In control (n = 3), Kir4.1 wt (n = 6) and Kir4.1 ko (n = 5) cells peak amplitude, rise time and decay time of mPSPs did not change after adding the GABA_A_R blocker, picrotoxin (150 µM). D) The inter event interval of mPSPs in Kir4.1 wt cells decreased in the presence of picrotoxin (* p = 0.02). Mann-Whitney U test; see Suppl. Table 2 for further details.

**Suppl. Figure 4:**
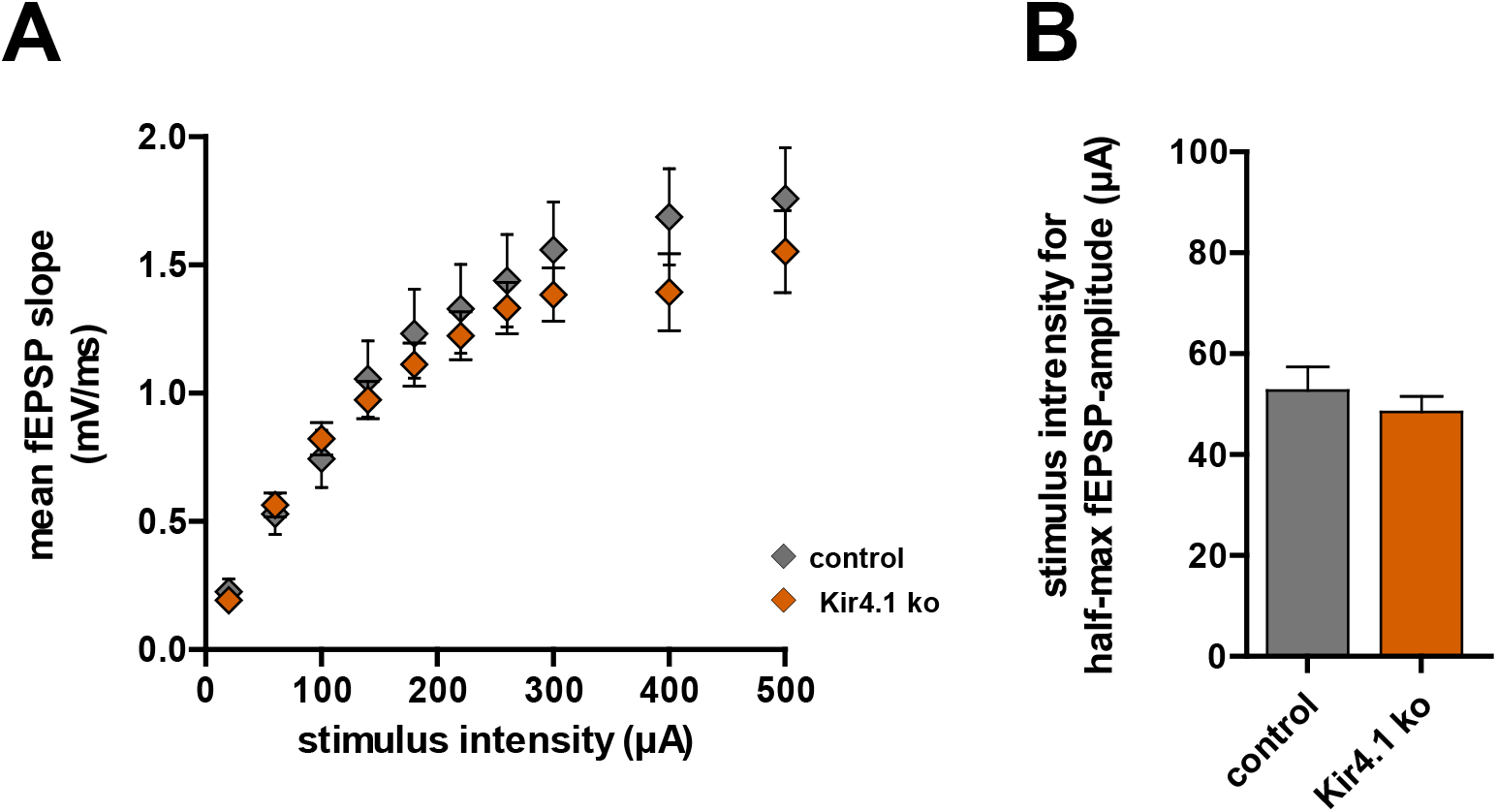
Basal excitability in hippocampal CA1 stratum radiatum is similar for slices of control and Kir4.1 ko mice. A) Input-output curves of slices from control (n=13; N=4) and Kir4.1 ko mice (n=35; N=12) show fEPSP slopes as a function of stimulus intensity in the CA1 stratum radiatum. B) Quantification of half-maximum fEPSP amplitude evoked by different stimulus intensities revealed similar sensitivity to electrical stimulation between genotypes. T-test, p > 0.05.

**Suppl. Table 1.**
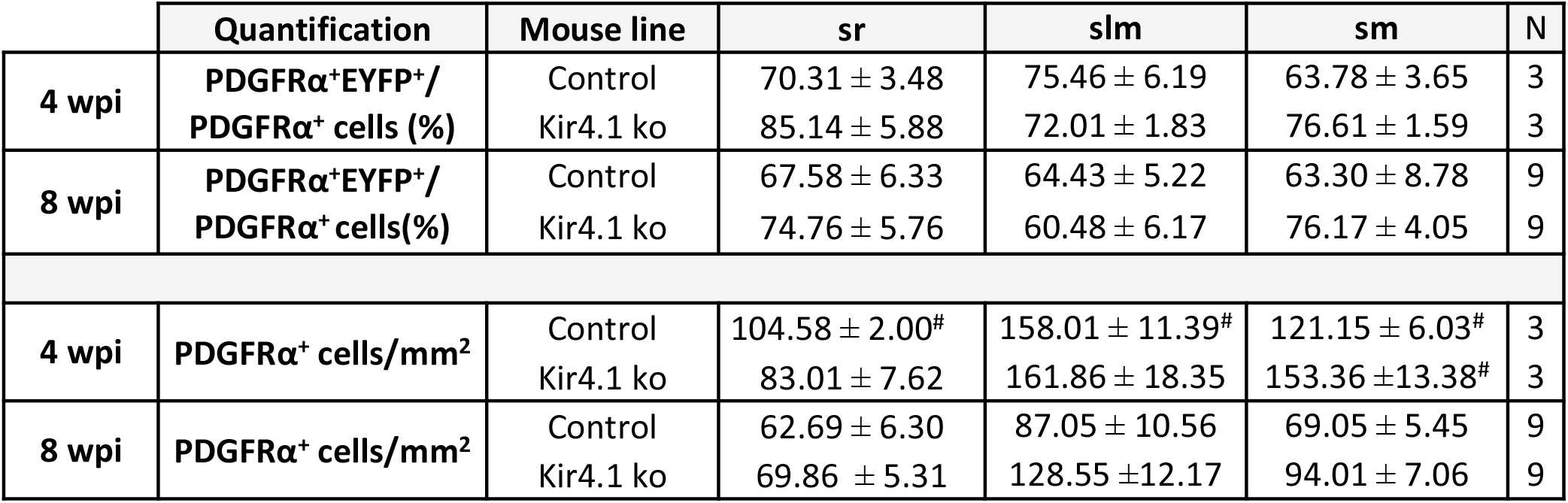
Recombination efficiency and density of hippocampal NG2 glia in control and Kir4.1 ko mice. Recombination efficiency was calculated by determining the number of cells being positive for EYFP and PDGFRα (EYFP^+^PDGRa^+^ cells) among all PDGRa^+^ cells, 4 and 8 weeks post tamoxifen injection. Data are given as mean ± SEM. N, number of mice; wpi, weeks post injection. sr, stratum radiatum; slm, stratum lacunosum moleculare; sm, stratum moleculare of the dentate gyrus. **^#^** Indicates significant differences between different time points (4 vs. 8 wpi) within the same genotype.

|  | Quantification | Mouse line | sr | slm | sm | N |
| --- | --- | --- | --- | --- | --- | --- |
| 4 wpi | PDGFRα<sup>+</sup>EYFP<sup>+</sup>/<br>PDGFRα<sup>+</sup> cells (%) | Control | 70.31 ± 3.48 | 75.46 ± 6.19 | 63.78 ± 3.65 | 3 |
|  |  | Kir4.1 ko | 85.14 ± 5.88 | 72.01 ± 1.83 | 76.61 ± 1.59 | 3 |
| 8 wpi | PDGFRα<sup>+</sup>EYFP<sup>+</sup>/<br>PDGFRα<sup>+</sup> cells(%) | Control | 67.58 ± 6.33 | 64.43 ± 5.22 | 63.30 ± 8.78 | 9 |
|  |  | Kir4.1 ko | 74.76 ± 5.76 | 60.48 ± 6.17 | 76.17 ± 4.05 | 9 |
| 4 wpi | PDGFRα<sup>+</sup> cells/mm<sup>2</sup> | Control | 104.58 ± 2.00<sup>#</sup> | 158.01 ± 11.39<sup>#</sup> | 121.15 ± 6.03<sup>#</sup> | 3 |
|  |  | Kir4.1 ko | 83.01 ± 7.62 | 161.86 ± 18.35 | 153.36 ± 13.38<sup>#</sup> | 3 |
| 8 wpi | PDGFRα<sup>+</sup> cells/mm<sup>2</sup> | Control | 62.69 ± 6.30 | 87.05 ± 10.56 | 69.05 ± 5.45 | 9 |
|  |  | Kir4.1 ko | 69.86 ± 5.31 | 128.55 ± 12.17 | 94.01 ± 7.06 | 9 |

**Suppl. Table 2.**
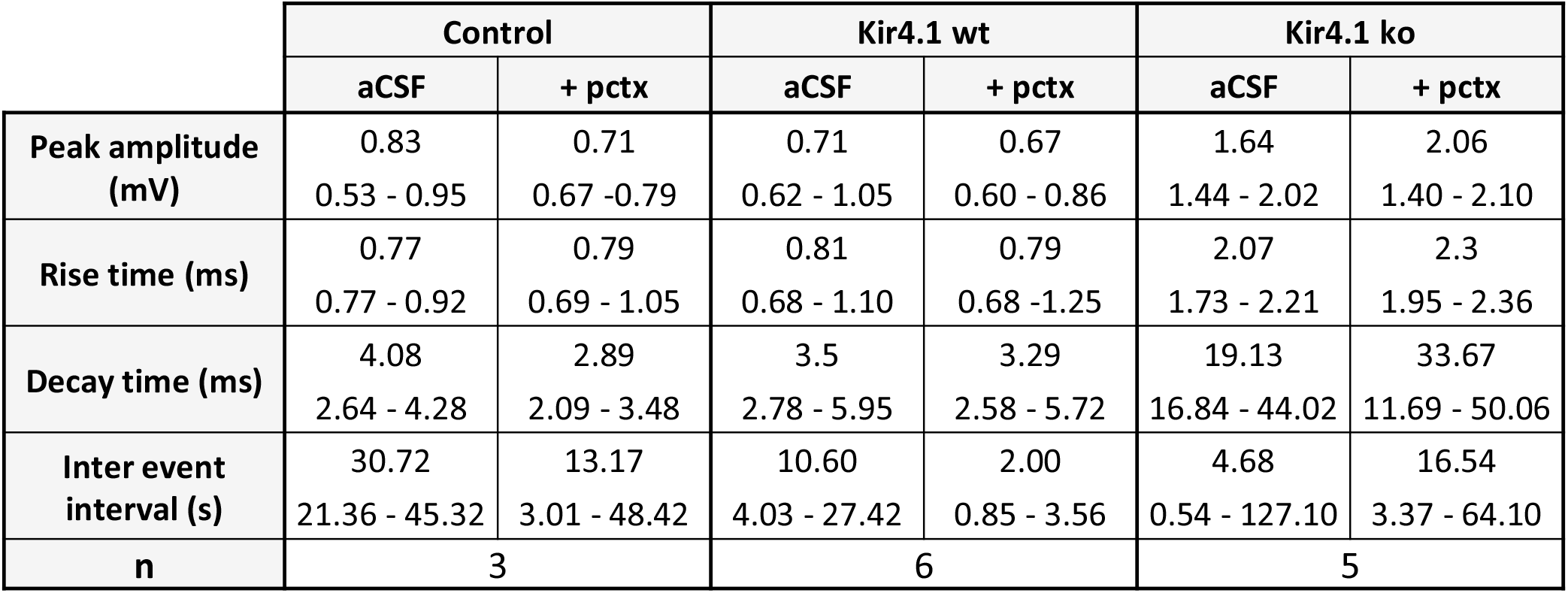
Characteristics of miniature EPSPs in control, Kir4.1*wt* and Kir4.1ko cells before and after application of picrotoxin (150 µM). Data is given as median and interquartile range (quartile 25%- quartile 75%). n, number of cells. pctx, picrotoxin. Mann-Whitey U Test.

|  | Control |  | Kir4.1 wt |  | Kir4.1 ko |  |
| --- | --- | --- | --- | --- | --- | --- |
|  | aCSF | + pctx | aCSF | + pctx | aCSF | + pctx |
| **Peak amplitude (mV)** | 0.83<br>0.53 - 0.95 | 0.71<br>0.67 - 0.79 | 0.71<br>0.62 - 1.05 | 0.67<br>0.60 - 0.86 | 1.64<br>1.44 - 2.02 | 2.06<br>1.40 - 2.10 |
| **Rise time (ms)** | 0.77<br>0.77 - 0.92 | 0.79<br>0.69 - 1.05 | 0.81<br>0.68 - 1.10 | 0.79<br>0.68 - 1.25 | 2.07<br>1.73 - 2.21 | 2.3<br>1.95 - 2.36 |
| **Decay time (ms)** | 4.08<br>2.64 - 4.28 | 2.89<br>2.09 - 3.48 | 3.5<br>2.78 - 5.95 | 3.29<br>2.58 - 5.72 | 19.13<br>16.84 - 44.02 | 33.67<br>11.69 - 50.06 |
| **Inter event interval (s)** | 30.72<br>21.36 - 45.32 | 13.17<br>3.01 - 48.42 | 10.60<br>4.03 - 27.42 | 2.00<br>0.85 - 3.56 | 4.68<br>0.54 - 127.10 | 16.54<br>3.37 - 64.10 |
| **n** | 3 |  | 6 |  | 5 |  |

**Suppl. Table 3.**
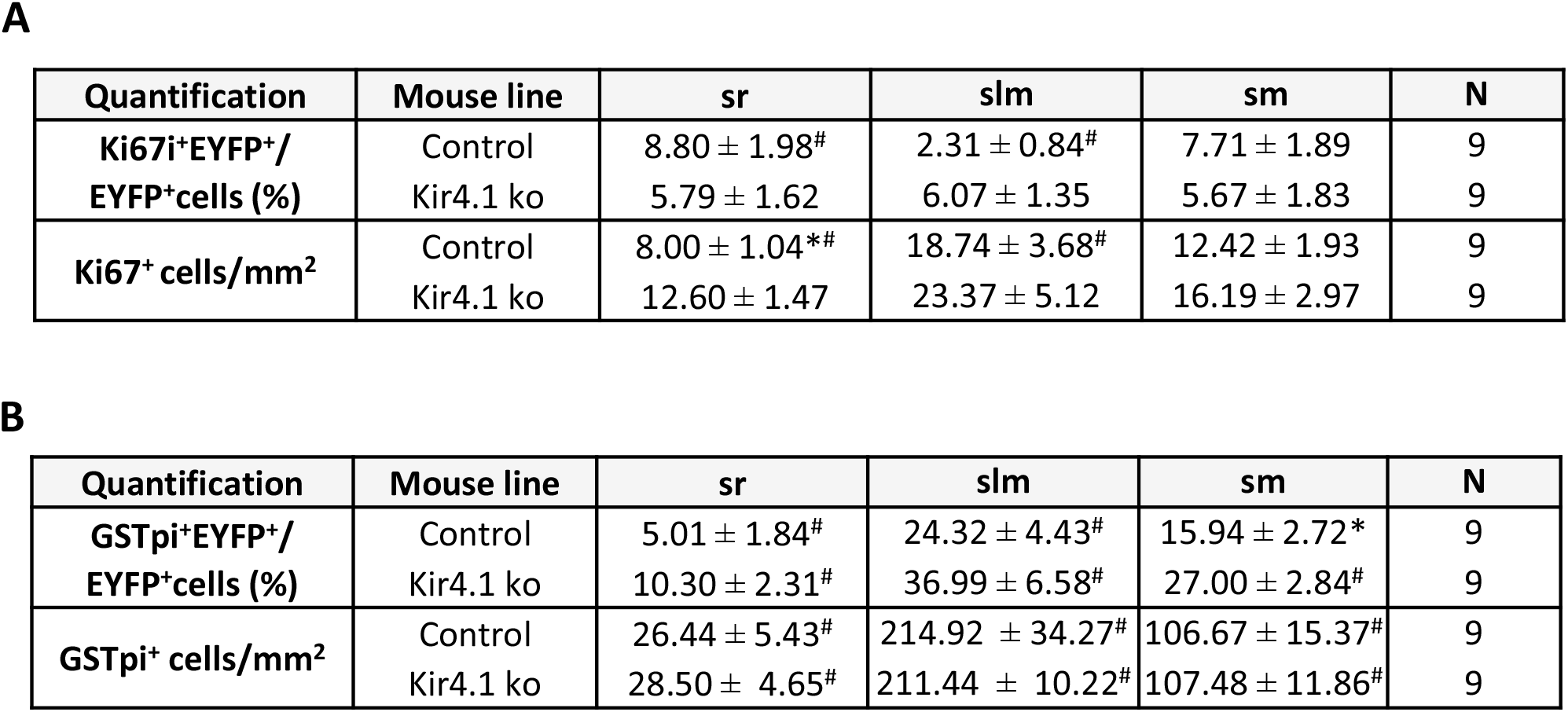
Proliferation and differentiation of hippocampal NG2 glia in control and Kir4.1 ko mice. A) Proliferative activity of recombined NG2 glia was calculated by determining the number of cells being positive for EYFP and Ki67 (EYFP^+^Ki67^+^ cells) among all EYFP^+^ cells. B) The number of recombined NG2 glia differentiating into oligodendrocytes was calculated by determining the number of cells being positive for EYFP and GSTpi (EYFP^+^GSTpi^+^ cells) among all EYFP^+^ cells. Data give mean ± SEM. N, number of mice. sr, stratum radiatum; slm, stratum lacunosum moleculare; sm, stratum moleculare of the dentate gyrus. Significant differences between genotypes within the same region (control vs. Kir4.1 ko mice) are indicated with **\***, between different subregions within the same genotype with ^#^.

| Quantification | Mouse line | sr | slm | sm | N |
| --- | --- | --- | --- | --- | --- |
| Ki67i+EYFP+/<br>EYFP+cells (%) | Control | 8.80 ± 1.98<sup>#</sup> | 2.31 ± 0.84<sup>#</sup> | 7.71 ± 1.89 | 9 |
|  | Kir4.1 ko | 5.79 ± 1.62 | 6.07 ± 1.35 | 5.67 ± 1.83 | 9 |
| Ki67+ cells/mm<sup>2</sup> | Control | 8.00 ± 1.04<sup>*#</sup> | 18.74 ± 3.68<sup>#</sup> | 12.42 ± 1.93 | 9 |
|  | Kir4.1 ko | 12.60 ± 1.47 | 23.37 ± 5.12 | 16.19 ± 2.97 | 9 |

**Suppl. Table 3. Proliferation and differentiation of hippocampal NG2 glia in control and Kir4.1 ko mice.**
| Quantification | Mouse line | sr | slm | sm | N |
| --- | --- | --- | --- | --- | --- |
| GSTpi+EYFP+/<br>EYFP+cells (%) | Control | 5.01 ± 1.84<sup>#</sup> | 24.32 ± 4.43<sup>#</sup> | 15.94 ± 2.72<sup>*</sup> | 9 |
|  | Kir4.1 ko | 10.30 ± 2.31<sup>#</sup> | 36.99 ± 6.58<sup>#</sup> | 27.00 ± 2.84<sup>#</sup> | 9 |
| GSTpi+ cells/mm<sup>2</sup> | Control | 26.44 ± 5.43<sup>#</sup> | 214.92 ± 34.27<sup>#</sup> | 106.67 ± 15.37<sup>#</sup> | 9 |
|  | Kir4.1 ko | 28.50 ± 4.65<sup>#</sup> | 211.44 ± 10.22<sup>#</sup> | 107.48 ± 11.86<sup>#</sup> | 9 |

